# Dextromethorphan inhibits collagen transport in the endoplasmic reticulum eliciting an anti-fibrotic response in *ex-vivo* and *in vitro* models of pulmonary fibrosis

**DOI:** 10.1101/2023.04.19.537530

**Authors:** Muzamil M Khan, Joanna Zukowska, Juan Jung, George Galea, Nadine Tuechler, Aliaksandr Halavatyi, Christian Tischer, Per Haberkant, Frank Stein, Ferris Jung, Jonathan Landry, Arif M. Khan, Viola Oorschot, Isabelle Becher, Beate Neumann, Thomas Muley, Hauke Winter, Julia Duerr, Marcus A Mall, Mikhail Savitski, Rainer Pepperkok

**Affiliations:** Cell Biology and Biophysics Unit, European Molecular Biology Laboratory, Heidelberg, Germany; Translational Lung Research Center Heidelberg (TLRC), German Center for Lung Research (DZL), 69120 Heidelberg, Germany; Advanced light microscopy facility, European Molecular Biology Laboratory, Heidelberg, Germany; Proteomics Core Facility, European Molecular Biology Laboratory, Heidelberg, Germany; Genomics Core Facility, European Molecular Biology Laboratory, Heidelberg, Germany; Biobank Thoraxklinik, University Hospital Heidelberg, Heidelberg, Germany; Department of Pediatric Respiratory medicine, Immunology and Critical Care Medicine, Charité-Universitätsmedizin Berlin, Germany; Electron Microscopy Core Facility, European Molecular Biology Laboratory, Heidelberg, Germany; Centre for Bioimage Analysis, European Molecular Biology Laboratory, Heidelberg, Germany; Chemical Biology Core Facility, European Molecular Biology Laboratory, Heidelberg, Germany; Molecular Medicine Partnership Unit, European Molecular Biology Laboratory and Heidelberg University, Heidelberg, Germany; Institute for Computational Biomedicine (ICB), Faculty of Medicine, Heidelberg University and Heidelberg University Hospital, Heidelberg, Germany; German Center for Lung Research (DZL), Associated Partner Site, Berlin, Germany; Berlin Institute of Health at Charité-Universitätsmedizin Berlin, Berlin, Germany; Genome Biology Unit, European Molecular Biology Laboratory, Heidelberg, Germany

## Abstract

Excessive deposition of fibrillar collagen in the interstitial extracellular matrix (ECM) of human lung tissue causes fibrosis, which can ultimately lead to organ failure. Despite our understanding of the molecular mechanisms underlying the disease, a cure for pulmonary fibrosis has not yet been found. In this study, we screened an FDA-approved drug library containing 712 drugs and found that Dextromethorphan (DXM), a cough expectorant, significantly reduces the amount of excess fibrillar collagen deposited in the ECM in *in-vitro* cultured primary human lung fibroblasts (NHLF) and *ex-vivo* cultured human precision-cut lung slice (hPCLS) models of lung fibrosis. Reduced extracellular fibrillar collagen levels in the ECM upon DXM treatment are due to a reversible trafficking inhibition of collagen type I (COL1) in the endoplasmic reticulum (ER) in TANGO1 and HSP47 positive structures. Mass spectrometric analysis shows that DXM causes hyper-hydroxylation of proline and lysine residues on Collagen (COL1, COL3, COL4, COL5, COL7, COL12) and Latent-transforming growth factor beta-binding protein (LTBP1 and LTBP2) peptides coinciding with their secretion block. In addition, thermal proteome profiling of cells treated with DXM shows increased thermal stability of prolyl-hydroxylases such as P3H2, P3H3, P3H4, P4HA1 and P4HA2, suggesting a change in activity. Transcriptome analysis of pro-fibrotic stimulated NHLFs and hPCLS upon DXM treatment showed activation of an anti-fibrotic program via regulation of pathways such as those involved in the MMP-ADAMTS axis, WNT, and fibroblast-to-myofibroblast differentiation. Taken together, the data obtained from both in-vitro and ex-vivo models of fibrogenesis show that Dextromethorphan has potent anti-fibrotic activity by efficient inhibition of COL1 membrane trafficking in the ER.

## Introduction

Fibrosis is defined as aberrant excessive deposition (also known as scarring) of extracellular matrix (ECM) in a given tissue^1^. It affects various organs in the human body, lungs, liver, kidney, intestines, skin to name a few and diseases^1,2^, such as cancer^2^ (tumour formation and metastasis), ageing^3^, SARS-CoV2 infected lungs^4^ have fibrosis as a comorbidity that contributes to the ultimate demise of the patients. In human lung pathology, the term Interstitial Lung Disease (ILD) is used for a group of diseases that causes fibrosis (or scarring). It is a result of persistent injuries to cells around alveoli that leads to heightened inflammatory tissue signalling and fibrotic scarring of lung tissue^1^. Fibrotic scarring causes tissue stiffness, which causes breathing difficulties and reduces oxygen levels in the bloodstream. As of 2019, in WHO European region, 761,000 people are suffering from ILD, 25,000 patients have died (in 2019) due to ILD, and 496,000 healthy years have been lost due to patients suffering from ILD^5^.

Currently, two drugs, Pirfenidone and Nintedanib, are FDA-approved for use against lung fibrosis^6^. However, these drugs do not cure the disease, only slow the rate of lung function decline, with patients responding variably to the drug treatment. In randomized clinical trials of both these drugs, 20-30% of patients permanently discontinued their treatment due to adverse side effects such as diarrhoea or photosensitivity^7^. Moreover, due to their high prices, these drugs have reduced availability. Therefore, to ameliorate or cure lung fibrosis, new drugs must be discovered that can reduce excess ECM in ILD pathology

So far as the molecular mechanism underlying pulmonary fibrosis is considered, it is known that in lung fibrosis patients, fibrosis occurs in the form of foci distributed in lung parenchyma^8^. These foci are created due to injury to lung alveoli, which results in inflammatory signalling involving activation pathways mediated by PDGF, TGFb1, Wingless/Int (WNT) and YAP/ TAZ signalling^9,10^. Among these, TGFb1 is considered to be the master regulator of lung fibrosis^11^ and fibrosis in general. The TGFb1 activation (together with other pathways) results in the trans-differentiation of interstitial fibroblast into myofibroblasts resistant to apoptosis^1^. Myofibroblasts are highly contractile cells expressing high levels of alpha-smooth muscle actin (α-SMA/ ACTA2) and synthesize copious amounts of ECM to be deposited in the interstitial space of lung parenchyma^1,8^. The persistent presence of cells (myofibroblasts) that can synthesis ECM at high rates results in the replacement of functional tissue (alveoli) with non-functional ECM (fibrotic foci)^1,8^. Therefore, lung fibrotic foci are characterized by abnormally high content of extracellular matrix (ECM) with an excess of fibrillar collagens (type I, III and V)^12,13^. At the ECM level, the excessive presence of these fibrillar collagens (primarily collagen type I) is the physical cause of eventual mortality^14^.

To this end, in this study, we have used ECM-deposited collagen type I (here onwards referred to as COL1) as a phenotypic marker in a high throughput microscopy-based screen of an FDA-approved library (Table-S8-DrugLibraryList). To screen for potential anti-fibrotic drugs, we optimized the so-called “scar-in-a-jar assay”^15,16^. Here, we investigated Dextromethorphan (among the top hits) for its potential anti-fibrotic activity. Anti-fibrotic activity and potential mechanism of action of Dextromethorphan were evaluated in *in-vitro* cultured primary fibroblast and ex-vivo cultured human precision-cut lung slice models of fibrosis (Figure 1D).

**Figure 1.**
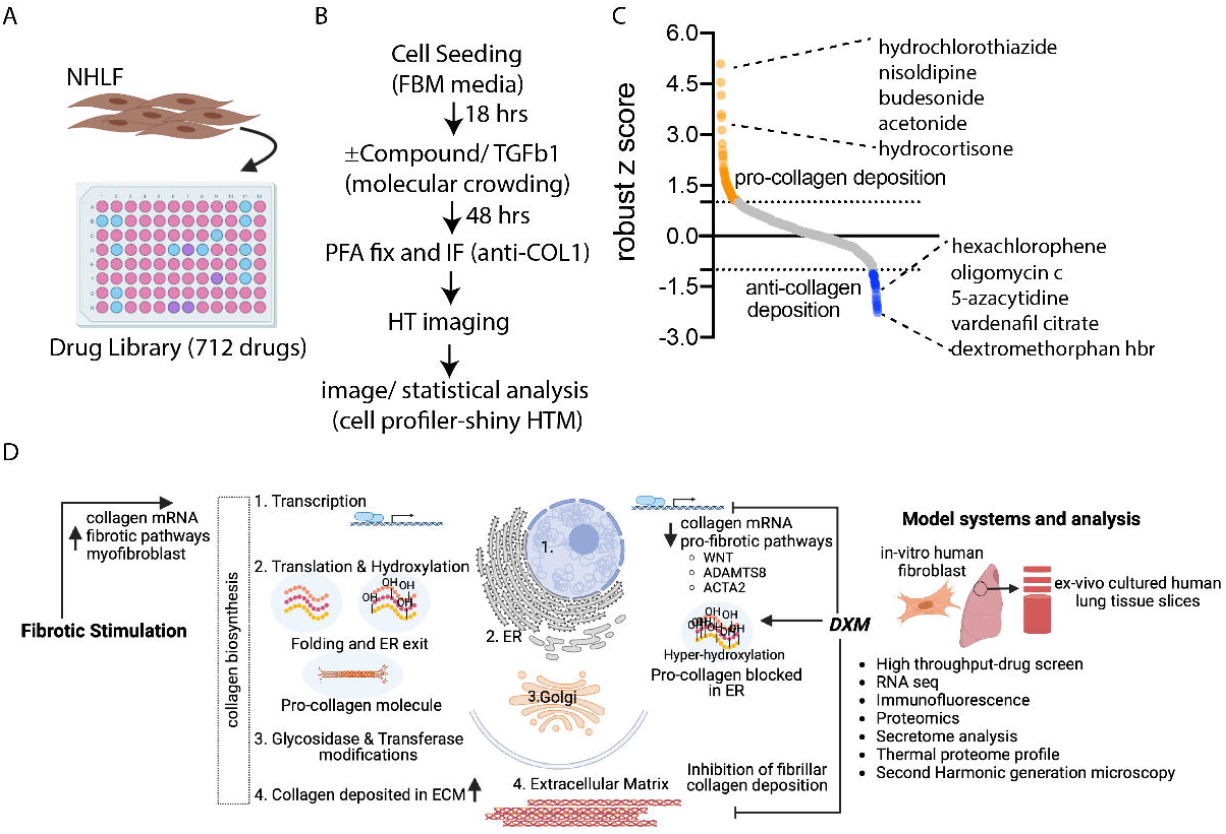
High-throughput screening and image analysis of FDA approved drug library reveals Dextromethorphan as a potential anti-lung fibrosis drug candidate. A-B, Schematic representation of high throughput screen set-up. C, robust z-scores of the respective drugs calculated using automated image analysis pipeline in cell profiler and shinyHTM. Graph shows top 5 drugs that showed a pro- and anti-collagen deposition effect respectively. D, Schematic representation of workflow used in the study, summarizing potential mechanism of anti-fibrotic action mediated by Dextromethorphan (DXM) in different models of lung fibrosis. Up and downward arrows represent upregulation and downregulation of respective terms. Blunted arrow heads (--|) represent inhibition and sharp arrow heads (➔) represent induction. Collagen mRNA after transcription is translated into ER, where it undergoes post-translational modification such as hydroxylation, glycosylation before passing via Golgi complex to Extracellular matrix for deposition. Upon fibrotic stimulation (e.g. TGFb1 treatment), collagen transcription is upregulated. In summary, DXM treatment causes hyper-hydroxylation of proline and lysine residues of collagen polypeptides, resulting in trafficking inhibition from Endoplasmic reticulum (ER). Accumulation of collagen protein in ER in a feedback signalling cascade suppresses expression of pro-fibrotic pathways at the mRNA level.

## Results

### FDA-approved drug library screen reveals Dextromethorphan as a potential anti-pulmonary fibrosis drug

A drug library consisting of 712 FDA-approved drugs was screened for inhibitors of ECM-deposited collagen type I [COL1] (Figure 1 A-C). Primary normal human lung fibroblasts (NHLF) were stimulated with TGFb1 in the presence of macromolecular crowding (MMC, Figure S1) to induce excess collagen synthesis and deposition in extracellular matrix. 48 hours post TGFb1 stimulation and drug treatment, deposition of COL1 in ECM was quantified by immunostaining and automated quantitative microscopy (Fig.1). Both positive controls i.e., Nintedanib (robust z-score of −0.53 and −0.98) and Pirfenidone (robust z-score of −0.57 and −0.93) reduced the deposition of COL1 in ECM (Table-S9-DrugRobustZScores). The median robust z-score of Nintedanib (−0.87) was taken as the threshold to select for the most efficient anti-fibrotic drugs. This classification revealed 5 potential anti-fibrotic drugs, which showed increased inhibition of COL1 deposition compared to the positive controls (Figure 1C). These (see Table-S8-DrugLibraryList) included two drugs with disinfectant properties (Hexachlorophene with robust z-scores of −2.19 and −2.42, and Oligomycin C with robust z-scores of −2.24 and −1.55). Two other drugs with known anti-fibrotic activity^17,18^, Vardenafil (robust z-scores of −0.38 and −2.59) and 5-azacytidine (robust z-score of −1.35 and −2.02) have earlier been shown not to be clinically efficient^17–19^. Therefore, among the top hits that reduced ECM-deposited COL1, d-3-methoxy-n-methylmorphinan hydrobromide (robust z-score of −1.17 and - 1.51) also known as Dextromethorphan Hbr, was chosen and further investigated in more detail as a potential novel anti-pulmonary fibrosis drug. Dextromethorphan (DXM) is approved as an antitussive drug (against cough)^20^. In cough suppressants, Dextromethorphan inhibits the NMDA receptor mediated neuronal signalling that evokes coughing reflex^20^.

Consistent with our screening data, we validated the effect of DXM on COL1 deposition with DXM compounds obtained from a commercial source different from the one used in the screening experiments (Figure 2A). Furthermore, a significant inhibition of ECM-deposited COL1 after DXM treatment was also observed in different human cells originating from lung, skin or kidney (Figure 2B). Next, we validated the effect on COL1 deposition in ECM using label-free Second Harmonic Generation (SHG) microscopy^21^, which allows the quantitative detection of fibrillar collagen (Figure 2C-D). Here, NHLF were treated with TGFb1 and DXM for one week. Acquired images and subsequent image analysis show and confirm a significant decrease of fibrillar collagen in ECM of NHLFs cultures upon concomitant treatment of DXM and TGFb1 (Figure 2D). Furthermore, titration of different DXM concentrations (Figure 2E) shows 10µM DXM as the minimum effective concentration inhibiting COL1 deposition.

**Figure 2.**
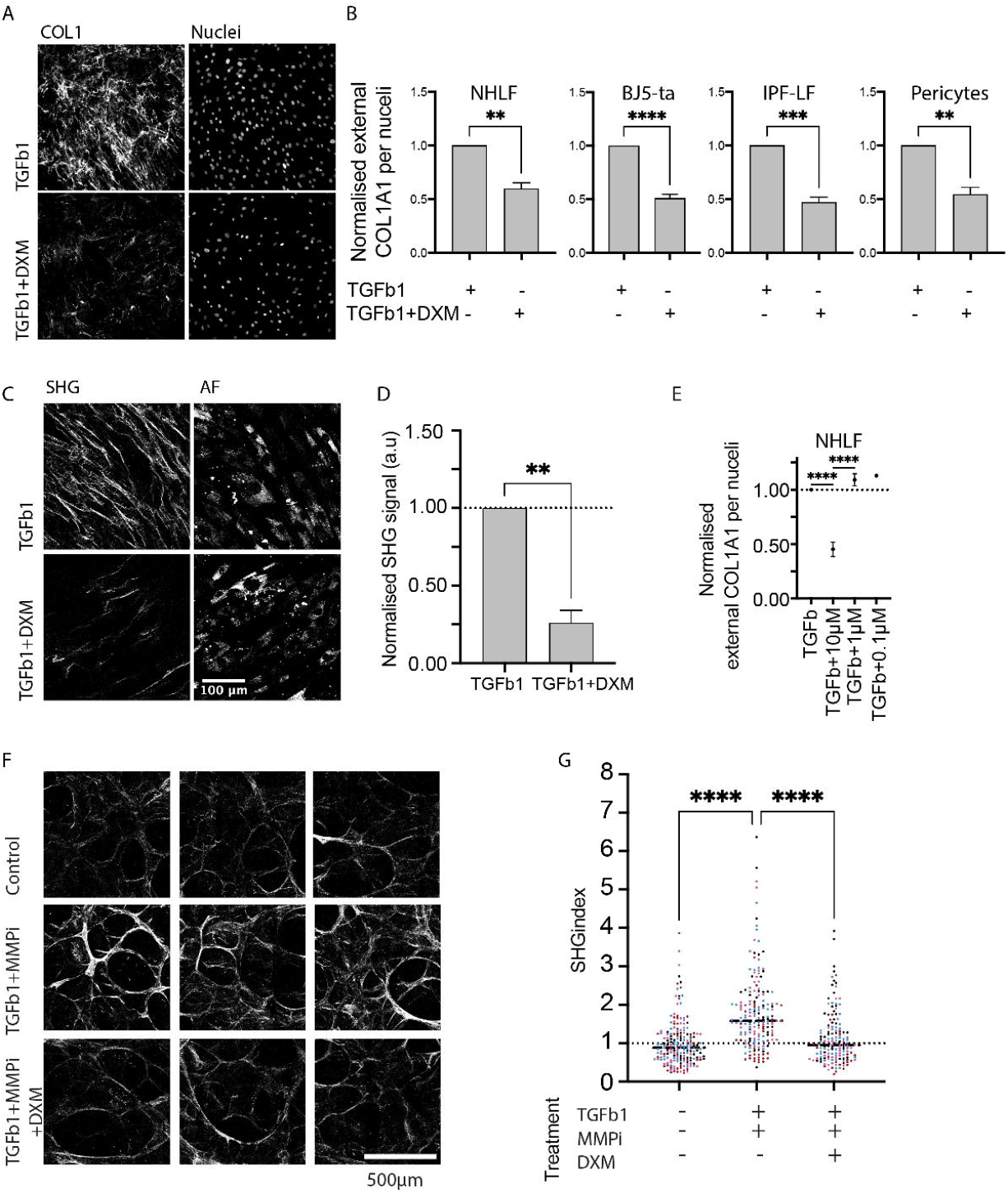
Dextromethorphan (DXM) inhibits excess deposition of fibrillar collagen in ex-vivo and in-vitro model systems of fibrogenesis. A, immunofluorescence staining against ECM deposited COL1 in NHLFs upon TGFb1 and TGFb1+DXM treatment. DXM used in this assay was obtained from a different commercial source. B, quantification of ECM deposited COL1 (immunofluorescence) by different cell types upon *in-vitro* culture. Normal Human Lung Fibroblasts (NHLFs), BJ-5ta, hTERT-immortalized foreskin fibroblasts, IPF-LF, Lung fibroblasts obtained from idiopathic pulmonary fibrosis patient, Pericytes, immortalized kidney pericytes. C, label-free SHG imaging of ECM deposited fibrillar collagen by NHLFs after 7 days of in-vitro culture (TGFb1 and TGFb1+DXM). SHG images represent fibrillar collagen signal, while AF represents 2-photon excited autofluorescence that gives an estimate of number of cells in the image field of view. D, quantification of SHG signal in each image normalised to AF, with subsequent normalization to the SHG signal from TGFb1 stimulated NHLFs. E, quantification of varying concentration of DXM on ECM deposited COL1 (immunofluorescence) in *in-vitro* cultured NHLFs. F, representative images (sum projections) of regions of interest from label-free SHG signal acquired from hPCLS (5mm in diameter and 350µm thickness). The hPCLS were stimulated with DMSO, TGFb1+MMPi and TGFb1+MMPi+DXM. Scale bar = 500µm. G, quantification of SHG signal across respective conditions. Color of the dot represents patient ID. Each dot represents a ROI analysed in respective hPCLS of the patient. 3-4 hPCLS were imaged per patient per condition. Patient n= 4. One-way ANOVA was used for testing statistical significance. p-value ≤0.001****. Note: FDA screen image analysis was performed on two replicates of experiments, while all other analysis in the figure here had n=3. Students t-test was used for testing statistical significance. p-value ≤0.01**, p-value ≤0.001***.

To analyze the efficacy of DXM on fibrillar collagen deposition in a more physiological fibrogenesis model, we used our previously established human precision-cut lung slice (hPCLS) culture model system^21^. hPCLSs were treated with TGFb1+MMPi to induce pro-fibrotic deposition of fibrillar collagen for two weeks (Figure 2F) and deposition of fibrillar collagen was quantified by label-free SHG imaging (Figure 2G). Consistent with our observations in NHLF cultures (Figure 2A-D), DXM treatment significantly reduced fibrillar collagen deposition in the ECM compared to TGFb1+MMPi control hPCLSs (Figure 2F-G).

### DXM blocks collagen type 1 and like cargoes in the endoplasmic reticulum

To gain further insights into the anti-fibrotic molecular mechanism of action of DXM, we performed proteomic analysis of NHLFs upon treatment with DMSO, TGFb1 and TGFb1+DXM. In total 5899 proteins were detected (Figure 3A). Comparison of TGFb1 and DMSO treated NHLFs showed that in total 150 proteins significantly (log2FC ≥ −0.58 or ≤ 0.58, p.adj. 0.05) changed their abundance upon TGFb1 treatment (Figure 3A). Pathway enrichment analysis of these significantly regulated proteins showed enrichment of pro-fibrotic pathways^22,23^ such as ECM organisation, Cell Adhesion and Tissue morphogenesis (Figure 3B). TGFb1 treatment caused a significant upregulation of pro-fibrotic ECM proteins^24^ such as COL1, COL3, FN1, THBS1, FBN1 (Figure 3B) and an upregulation of enzymes that aid the polymerization of collagens in ECM such as BMP1, LOXL2, COL5^25^. To analyze how concomitant treatment with DXM and TGFb1 is regulating the pro-fibrotic proteins and pathways, we analysed the significantly regulated (log2FC ≥ −0.58 or ≤ 0.58, p.adj. 0.05) proteins and in particular the levels of proteins involved in ECM organisation (Figure 3C-D, TGFb1+DXM vs TGFb1). This analysis revealed significantly increased intracellular levels of ECM proteins COL1, COL5, COL7, COL12, COL15, COL16 and FN1 in TGFb1+DXM treated compared to NHLFs treated with TGFb1 alone (Figure 3D). One explanation for this result together with our observed inhibition of COL1 extracellular deposition could be that DXM induces an intracellular accumulation of COL1. In order to test this hypothesis, we localized intracellular COL1 by immuno-fluorescence. Confocal imaging revealed a prominent cytoplasmic accumulation of COL1 in DXM treated cells (Figure 3E, zoom, white arrow heads).

**Figure 3.**
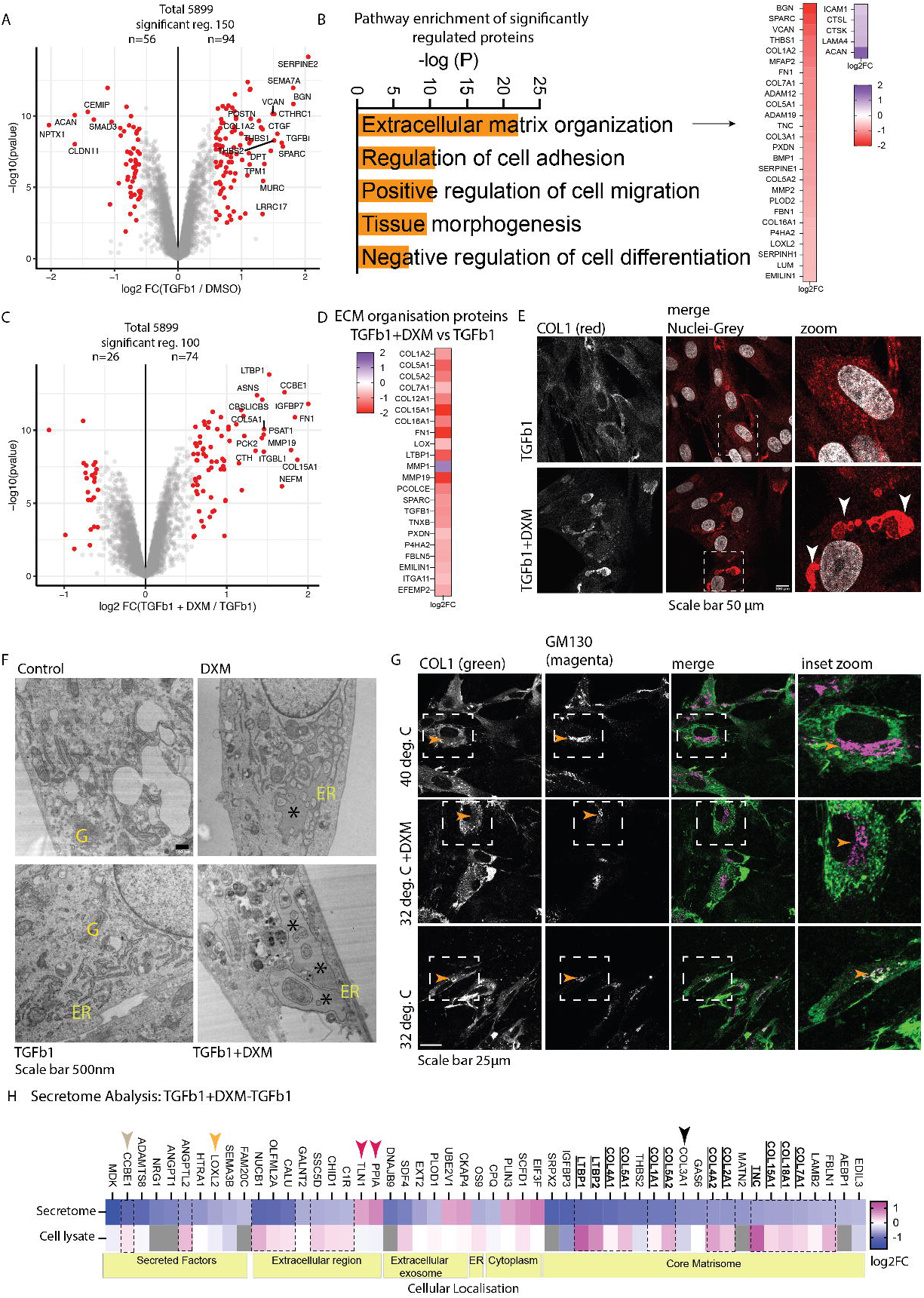
DXM induces collagen transport inhibition in endoplasmic reticulum. A, volcano plot analysis of differentially (TGFb1 vs DMSO) regulated proteins (mass-spectrometry) in NHLFs. Red dots represent the top hits, blue represents potential candidates and green represent proteins that did not pass the significance criteria. B, Pathway enrichment of significantly regulated (log2FC between 0.58 or −0.58, padj ≤ 0.05) proteins using Metascape gene ontology database. C, Volcan plot analysis of differentially (TGFb1 vs TGFb1+DXM) regulated proteins (mass-spectrometry) in NHLFs. D, Log2FC values of listed proteins (TGFb1 vs TGFb1+DXM). E, immunofluorescence staining of endogenous COL1 protein after permeabilization of in vitro cultured NHLFs (TGFb1 and TGFb1+DXM stimulated) (the images represent single optical plane from the respective z-stacks acquired using a confocal microscope). Red signal represents COL1 immunostaining and the nuclei (false colored in grey) were stained using Hoechst 33342. F, Transmission Electron Microscopy analysis of NHLFs cultured in vitro under different conditions (DMSO-Control, TGFb1, DXM or TGFB1+DXM stimulated). ER= Endoplasmic Reticulum, G= Golgi Complex. Scale bar IF image=50 µm, EM image=500 nm. G, collagen transport assay: immunofluorescence staining of COL1 in NHLFs treated with TGFb1 (24 hrs), incubated at 40°C to induce collagen trafficking ER block (for 6 hrs), and at 32°C (for 30 minutes) to allow release of COL1. Cells were also treated with DXM during the release (32 deg. C+DXM). Scale bar 25µm. H, secretome (MS analysis) column of heat map shows the log2FC of significantly regulated (log2FC between 0.58 or −0.58, padj ≤ 0.05) proteins in the media supernatant of NHLFs treated with TGFb1+DXM compared to TGFb1 treated NHLFs, Cell lysate column of the heat map shows the respective log2FC of proteins in the SDS solubilized cell lysates. Proteins undergoing trafficking inhibition are listed as bold and underlined. If the respective protein in secretome was not measured in cell lysates the heat map cells are colored grey. Black dashed rectangles highlight secreted proteins that show trafficking inhibition based reduced abundance in SNs. Black and yellow arrow heads show secreted protein that do not show trafficking based inhibition with corresponding under hydroxylated or no change in hydroxylation levels of amino acid residues.

To characterize the sub-cellular compartment of this DXM-induced COL1 transport inhibition, we performed co-immunostainings of COL1 (Figure S3) with ER (Calnexin), ER-Golgi intermediate compartment (ERGIC 53) and Golgi Complex (GM130) marker proteins. Strikingly, the DXM-induced COL1 accumulations did not localize to the Golgi complex (Figure S3A), the ER-Golgi intermediate compartment (Figure S3C), or Calnexin (Figure S3B, white arrow head). However, electron microscopy analysis of NHLFs treated with DMSO, TGFb1, DXM and TGFb1+DXM showed a bloated ER (Figure 3F, black asterisk) both upon DXM or TGFb1+DXM treatment, suggesting a DXM-induced block of collagen in the endoplasmic reticulum.

In order to investigate if this collagen block in the endoplasmic reticulum is due to transport inhibition in ER, TGFb1-treated NHLF were cultured at 40°C for 6 hours to arrest COL1 in the ER due to its inhibition in proper folding and prevent its transport to Golgi complex (Figure 3G) ^26^ (control, zoom inset, Figure 3G, 40°C). COL1 was transported to the Golgi complex upon incubation of cells at 32°C for 30 minutes (Figure 3G, 32°C, yellow arrow heads inset zoom). However, when the NHLFs were treated with DXM for 30 minutes at 32°C, COL1 transport to the Golgi complex was strongly inhibited in the ER (insert zoom, arrow head, Figure 3G, 32°C +DXM, white overlap region). These data demonstrate that the transport of COL1 is indeed inhibited in ER.

### Secretome analysis of NHLFs shows DXM induces trafficking inhibition of COL1 and like cargoes

To establish the specificity of DXM-induced trafficking inhibition, we performed mass spectrometry analysis of cell culture media supernatants (SNs) from NHLFs treated with TGFb1+DXM and TGFb1 alone (Figure 3H, Secretome column of heatmap). SDS soluble cell lysates (CLs) of the respective NHLFs were also analysed. Out of 742 proteins detected in SNs, 208 proteins have reviewed secretory annotation in Uniprot data base^27^ (Table-S11-Secretome-TGFb1+DXM-TGFb1-NHLF, Reviewed secreted proteins). Of these 208 proteins (Table-S11-Secretome-TGFb1+DXM-TGFb1-NHLF), only 38 showed a significant change (log2FC ≥ −0.58 or ≤ 0.58, p.adj. 0.05) in SNs of NHLFs treated with TGFb1+DXM compared to TGFb1 treatment alone. These proteins are classified under Secreted Factors, Extracellular region and Core matrisome compartments of cellular localization (Figure 3H, Cellular localization). 36 of these proteins had a reduced abundance (negative log2FC in Secretome column) and 2 had an increased abundance (positive log2FC in secretome column). 22 proteins (out of 38 proteins significantly changing proteins) had a decreased abundance in SNs and an increased abundance in CLs (dashed rectangle outline), suggesting a transport inhibition of the respective cargoes from inside of the cell to the extracellular space. To validate, that a decreased abundance in SNs with corresponding increased abundance in CLs is indeed due to trafficking block, we additionally analysed localisation of CCBE1 (Figure 3H, grey arrow head), a non-core matrisome secreted factor, via immunofluorescence upon TGFb1+DXM treatment. Indeed, consistent with mass spectrometric analysis, CCBE1 also showed COL1 like DXM-induced intracellular accumulation, suggesting a trafficking inhibition by DXM treatment (Figure S2A).

Core matrisome proteins that showed a trafficking inhibition included collagens [Figure3H, Secretome column of heatmap] (COL1A1, COL3A1, COL4A1, COL5A1/A2, COL7A1, COL15A1 and COL18A1), latent-transforming growth factor beta-binding protein 1 and 2 (LTBP1 and LTBP2), Fibulin 1 (FBLN1), and Tenascin (TNC). Furthermore, transforming growth factor beta-1 proprotein (TGFb1) also showed a reduced abundance in supernatants (log2FC of −0,76, fdr = 0.0505, Table-S11-Secretome-TGFb1+DXM-TGFb1-NHLF). LTBP1, LTBP2 and TGFb1 proproteins control the activity of TGFb1 elicited pathways^28^. These data suggest that in addition to reducing the deposition of collagens in ECM, DXM might also inhibit TGFb1 induced pro-fibrotic pathways. This potential dual action DXM could put significant halt to fibrosis progression. Furthermore, other matrisome associated proteins^29^ [Figure8A, Secretome column of heatmap] including PLOD1, HTRA1, LOXL2 (Figure 3H, Secretome column of heatmap) also showed a decrease abundance in media supernatants.

To confirm that the effect of DXM-induced transport inhibition occurs predominantly to collagens and like cargoes, we also performed a well-established Vesicular Stomatitis Virus G protein (VSVG) transport assay^30^ (Figure S1C-D). Previous studies have shown that collagen and VSVG are transported via independent COPII dependent pathways^31^.Here, NHLFs were infected with recombinant adenovirus expressing thermosensitive YFP-tagged tsO45G (VSVG protein)^30^ and subsequently incubated at 40°C for 18 hours, which results in a transport block of VSVG in the endoplasmic reticulum^30^ and prevents it from localizing to cell surface. Upon release of this transport block by incubating cells at 32°C, VSVG was transported through the secretory pathway to the plasma membrane (Figure S1C, Control). Treatment of cells with brefeldin A (Figure S1C, BFA^32^), a known inhibitor of protein transport from the endoplasmic reticulum to the Golgi complex, showed impaired transport of VSVG to the cell surface. In contrast, cells treated with DXM showed apparently comparable transport of VSVG protein to the cell surface as control cells (Figure S1C, DXM). Indeed, quantitative image analysis confirmed that DXM didn’t affect VSVG transport to plasma membrane (Figure S1D).

### DXM-induced collagen type 1 accumulations in ER are reversible and positive for TANGO and HSP47 in endoplasmic reticulum

Post-transcription, the nascent collagen chains are inserted into the ER lumen where they undergo hydroxylation (proline and lysine hydroxylation), glycosylation, and correct folding. The hydroxylation processes are carried out by a set of molecules such as Prolyl 3- and 4-hydroxylases (in complex with CYPB and CRTAP) and LH2-FKBP65^33^. For correct triple helical folding of COL1, PDI aids the formation of disulphide bonds and eventual folding of COL1 chains is looked over by chaperone protein HSP47^33^ (SERPINH1). Post folding into a procollagen triple helix, the triple helix is guided to ER exit sites where it is packed into large COPII vesicles with the help of a collagen transport specific protein TANGO1^34^ (MIA3) (for a detailed description of the process, please see the review Claeys et al., 2021, Human Genetics^33^). Immunofluorescence staining analysis of NHLFs treated with TGFb1 and TGFb1+DXM revealed that the COL1 accumulation in ER are positive for TANGO1 and HSP47 (Figure 4A-B). Furthermore, co-staining with protein disulphide isomerase (PDI) showed accumulation similar to COL1, albeit mutually exclusive (Figure S2B). ER exit-site marker, SEC31 didn’t not show any redistribution upon DXM treatment (Figure S2C).

**Figure 4.**
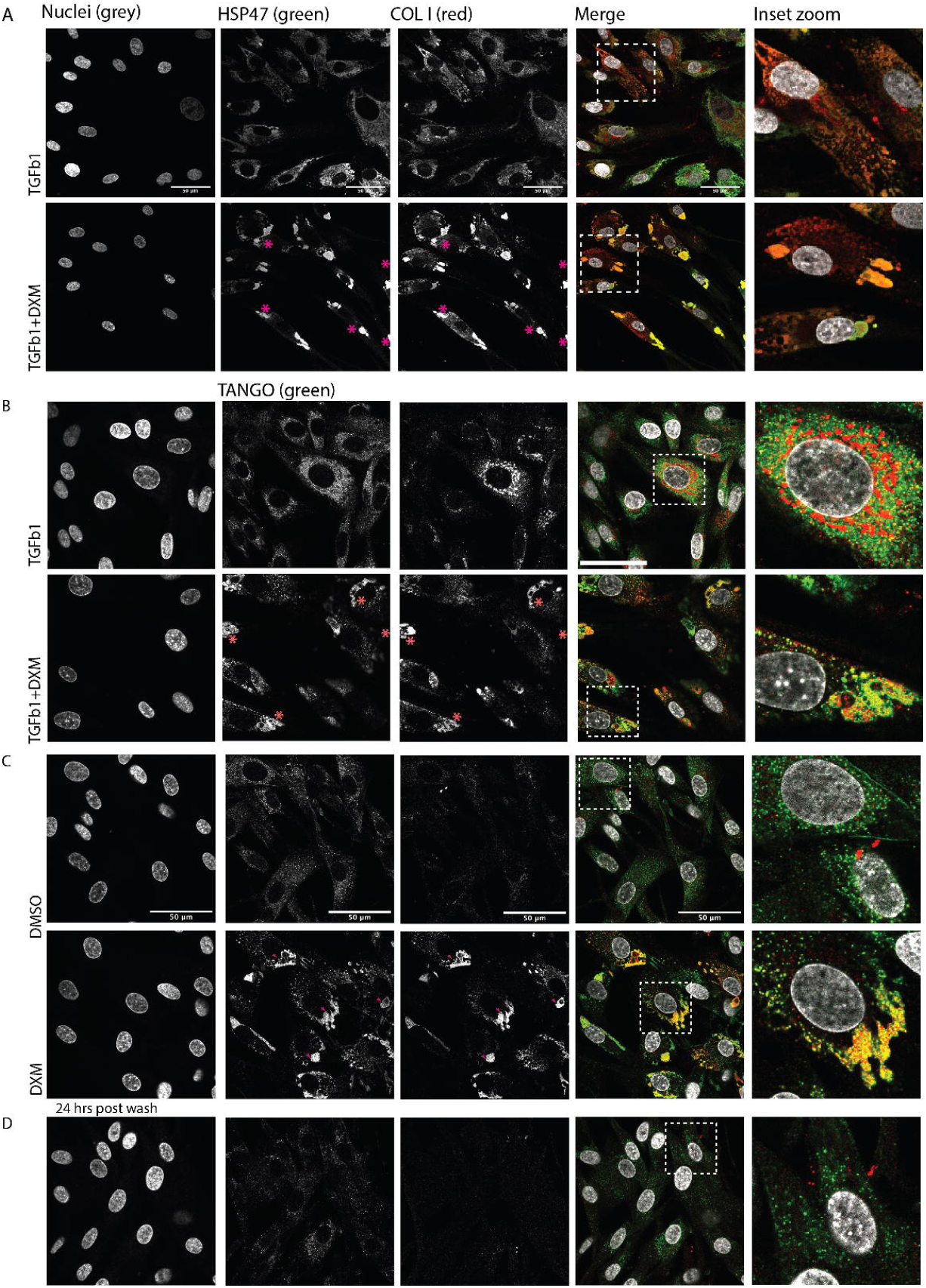
DXM causes a reversible redistribution of collagen transport specific proteins TANGO and HSP47. A**-**B, representative immunofluorescence staining acquired using a confocal microscope. The images represent single optical planes from the respective z-stacks. Nuclei were stained using Hoechst 33342, primary antibodies against HSP47, COL1 and TANGO (TANGO1) were used to visualize the respective endogenous proteins in NHLFs (TGFb1 and TGFb1+DXM stimulated). N=2 (number of time the experiment was repeated) for each immunostaining experiment. Scale bar= 50µm. C-D, representative confocal images of NHLF stained for COL1 and TANGO1 (TANGO) upon treatment with DMSO and DXM, B, after 24 hours wash out of the DXM drug. N=2. Inset zooms represent areas in dashed square boxes from merge column of images. Scale bar= 50µm.

To test if the DXM-induced accumulations of COL1 in endoplasmic reticulum are persistent after the removal of DXM, NHLFs were treated with DXM (for 48 hours). After 48 hours, DXM containing cell culture media was replenished with fresh media devoid of DXM for another 24hours. Next, immunofluorescence analysis of TANGO1 and COL1 in NHLFs treated with DMSO, DXM and 24-hour post wash of DXM was performed. Data analysis showed that both TANGO1 and COL1 distribution was indistinguishable from nontreated control cells (Figure 4C-D) suggesting that the DXM mediated COL1 accumulations in the ER are reversible at the time-scale of our experiments.

### Functional proteomics analysis of DXM treated NHLFs using thermal proteome profiling

Thermal proteome profiling (TPP)^35^ measures thermal stability of proteins on a proteome-wide scale and enables unbiased detection of protein ligand interactions due to the fact that ligand binding will typically increase thermal stability of proteins. Moreover, it has been shown that TPP can also detect thermal stability changes caused by changes in protein activity, protein-protein interactions and post-translational modifications^36,37^. In order to understand, how dextromethorphan could be inducing collagen specific trafficking inhibition, we performed TPP on NHLF treated with DSMO and DXM. To determine compound concentrations which might show stability of proteins that are specific to the phenotype of COL1 accumulation in ER, we treated NHLFs with various concentration (1, 2.5, 5 and 10µM, Figure 5A). Immunofluorescence analysis of internal collagen showed that DXM-induced COL1 accumulation at 5 and 10µM concentrations (yellow arrow heads, Figure 5A). These data provided the rationale for testing the thermal stability of proteins using TPP at 1, 5 and 10µM concentration of DXM treatment. Following this we looked at thermal stability changes of the 51 known proteins of Collagen 1 proteostasis network^38^ (Figure 5B). The data analysis showed that 21 proteins (bold italic, Figure 5B) showed increased thermal stability at 5 and 10µM DXM treatment (black dashed box, Figure 5B). Of note, these proteins include P3H2, P3H3, P3H4, P4HA1, P4HA2, prolyl-hydroxylase complex associated protein CRTAP and lysyl hydroxylase PLOD1 and PLOD2(Figure 5B, dashed black rectangle). These data suggest a potential higher activity of prolyl-hydroxylase enzymes. Furthermore, other chaperon proteins that are associated with collagen folding and were also stabilized include phosphodiesterase enzyme PDAI4, FKBP10 (also known as FKBP65), PCOLCE and HYOU1 [Figure 5B].

**Figure 5.**
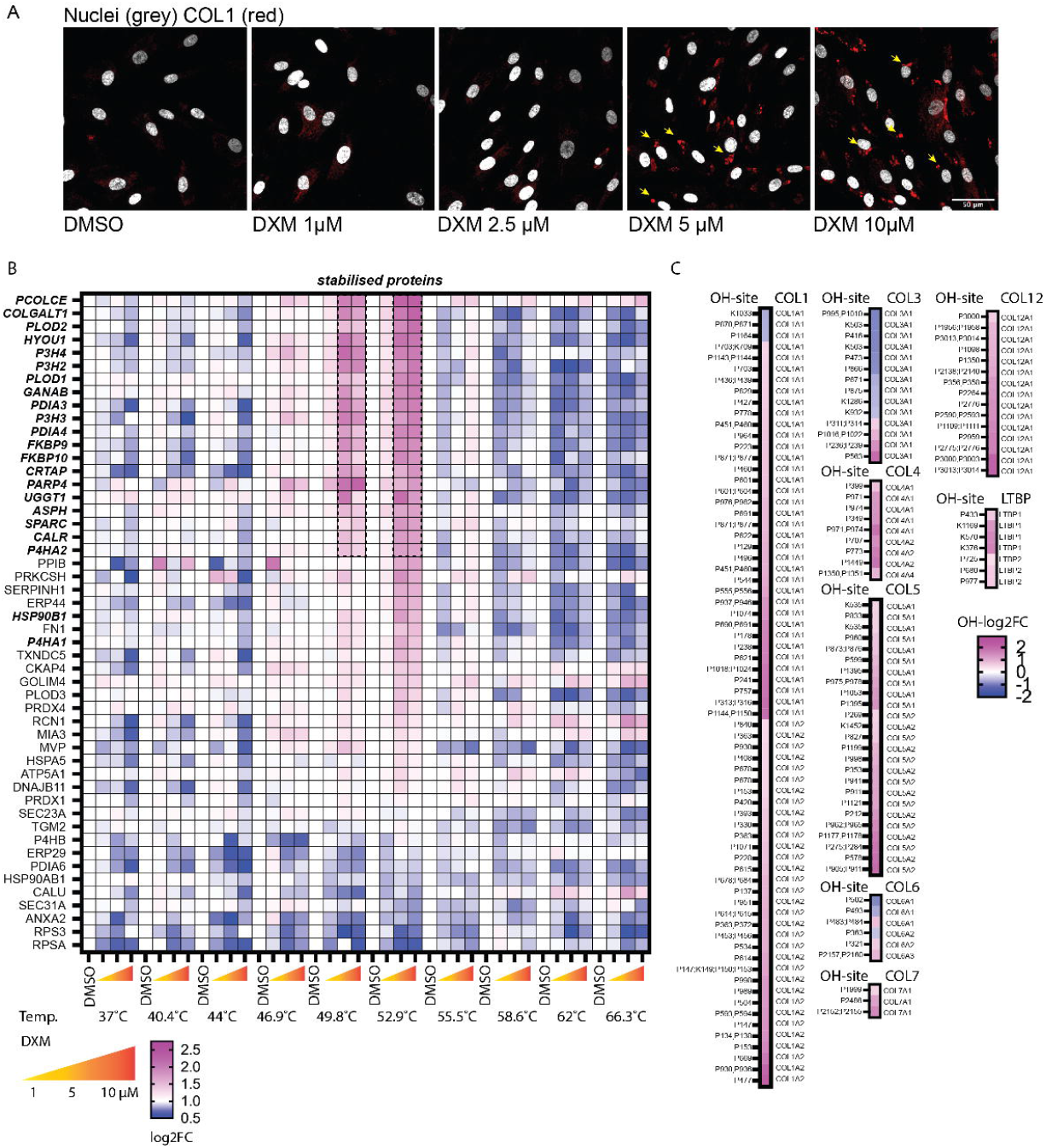
Thermal proteome profiling of DXM treated NHLFs show stabilization of proline and lysine hydroxylase enzymes involved in collagen proteostasis network: A, representative images of NHLFs treated with DMSO and respective concentration of DXM. Cells were stained for COL1. Yellow arrow heads show DXM induces intracellular accumulation of COL1 at 5 and 10µM concentration. B, heat-maps show the thermal stability of 51 known proteins that interact with COL1 across different DXM concentration treatments of NHLFs. Cell lysates from differentially treated NHLFs were heated at various temperatures (Temp) followed by MS analysis. Black dashed rectangle selection shows selective stabilization of proteins at 5 and 10µM DXM treatment. C, heatmap represents the log2FC changes in the hydroxylation levels of the respective proline and lysine residues of different collagens upon treatment with TGFb1+DXM compared to TGFb1.

### DXM treatment causes hyper-hydroxylation of Collagens in ER

Previous studies show that 2,2-dipyridil, a known hypoxia-inducible factor prolyl-hydroxylase inhibitor induces a block of collagen trafficking in ER with similar kinetics as observed in our model system^39^ here. In additional inhibition- or hyper activation of hydroxylase enzymes in osteogenesis imperfecta are known to cause collagen block in ER^40^. These studies, together with our results of TPP analysis that show stabilisation prolyl and lysyl hydroxylase enzymes raised the question, if hydroxylation levels of proline and lysine residues on proteins NHLFs treated with DXM are affected.

Furthermore, the relatively fast kinetics of the transport inhibition (Figure 3G, Figure S1 C-D) suggests changes in post-translational enzyme kinetics or post-translational modification of collagen protein in the ER. To this end, the peptides detected in mass spectrometry analysis of NHLF cell lysates (treated with TGFb1+DXM compared to TGFb1) were analysed for hydroxylation levels at various amino acids. The analysis revealed a total of 4289 peptides from 1455 proteins that were hydroxylated (Table-S12-OH-TGFb1+DXM-TGFb1-NHLF). Out of 4289 hydroxylated peptides, 231 peptides from 61 proteins were significantly (log2FC ≥ −0.58 or ≤ 0.58, p.adj. 0.05) changed upon TGFb1+DXM treatment compared to TGFb1-treated NHLFs (Table-S12-OH-TGFb1+DXM-TGFb1-NHLF, OH tab). Out of 231 (significantly changed), 185 peptides were significantly hyper hydroxylated and 83 peptides were significantly under hydroxylated (Table-S12-OH-TGFb1+DXM-TGFb1-NHLF, OH tab). 140 of the total (231) significant differentially hydroxylated peptides were collagens [COL1, COL3, COL4, COL5, COL6, COL7, COL12] (Figure 5C). Strikingly, all the collagens that showed significant hyper hydroxylation of proline and lysine residues (Figure 5C) were also transport inhibited (Figure 3H, bold underlined collagen names). Only COL3 and COL6 showed under hydroxylated residues. COL3 showed a reduced abundance in SNs but was also reduced in expression in CLs (Figure 3H, black arrow head). COL6 did not show a significant change in abundance in SNs (Table-S11-Secretome-TGFb1+DXM-TGFb1-NHLF, secretome tab). Furthermore,11 of 22 secreted proteins that showed a trafficking inhibition (Figure 3H, bold underlined) upon DXM treatment were also detected and measured in the hydroxylation analysis. Interestingly, all these 11 proteins (Figure 3H, bold underlined) had peptides that were significantly hyper hydroxylated (Table-S12-OH-TGFb1+DXM-TGFb1-NHLF, OH tab, yellow highlighted cells).

Notably, LOXL2 showed a significant decrease in abundance in SNs (Figure 3H, yellow arrow head) and also a decrease abundance in CLs, hence not a transport-based inhibition, showed no significant change in hydroxylation levels. Furthermore, 2 secreted proteins that showed an increased abundance in SNs, PPIA and TLN1 (Figure 3H, magenta arrow heads) did not show significant change in hydroxylation levels of proline and lysine amino acid residues (Table-S12-OH-TGFb1+DXM-TGFb1-NHLF). In addition, other secreted proteins that were significantly hydroxylated, were also hyper hydroxylated (Table-S12-OH-TGFb1+DXM-TGFb1-NHLF, OH-Secreted proteins tab). These proteins are FBN1, FN1, IGFBP7, MMP2, PCOLCE, PXDN and, SPARC. Notably, all these proteins (except FN1) showed a decreased abundance in SNs and increased abundance in CLs of NHLFs treated with TGFb1+DXM compared to TGFb1-treated NHLFs (Table-S11-Secretome-TGFb1+DXM-TGFb1-NHLF, Secretome and Cell lysate tab), but did not pass the significance threshold. These data suggest that hyper-hydroxylation of collagen and like cargoes could be the mechanism underlying the DXM-induced ER localised trafficking inhibition.

### Dextromethorphan induces transcriptional activation of an anti-fibrotic programme

Since DXM treatment caused a reduction in the abundance of TGFb1 activating proteins (LTBP1, LTBP2 and TGFb1 pro-protein) in the media supernatants of NHLFs, we investigated DXM caused a transcriptional inhibition of TGFb1 induced pro-fibrotic pathways. Here, we performed a mRNA seq analysis of TGFb1 and TGFb1+DXM treated NHLF cells and hPCLSs (Figure 6A-D). Data analysis of TGFb1-treated NHLF cells shows that markers of myofibroblasts such as alpha smooth muscle actin (*ACTA2*), Transgelin (*TAGLN*), Asporin (*ASPN*) are significantly upregulated with log2FC of 2.05, 0.81 and 2.24, respectively (Table-S2-Trancriptomics-NHLF). Pathway analysis (Figure 6A-B) of 1189 significantly (log2FC between 1 or −1, padj ≤ 0.05) regulated genes (Figure 6A) in TGFb1-treated NHLFs showed an increase in the expression of genes and pro-fibrotic pathways known to be regulated in fibroblasts (Figure 6A-B). Upregulated pathways included Naba core matrisome, blood vessel development, tube morphogenesis and integrin pathways, all known to be involved in fibrosis progression^41^. Gene enrichment of significantly down-regulated genes included inflammatory response, Naba matrisome associated signalling and regulation of epithelial cell proliferation among others (Figure 6B). Together, our ECM-deposited COL1 immunofluorescence data (Figure 2, Figure S1) and transcriptional profiling (Figure 6A-B) confirm a robust pro-fibrotic signalling environment present in the TGFb1-treated NHLFs used in this study.

**Figure 6.**
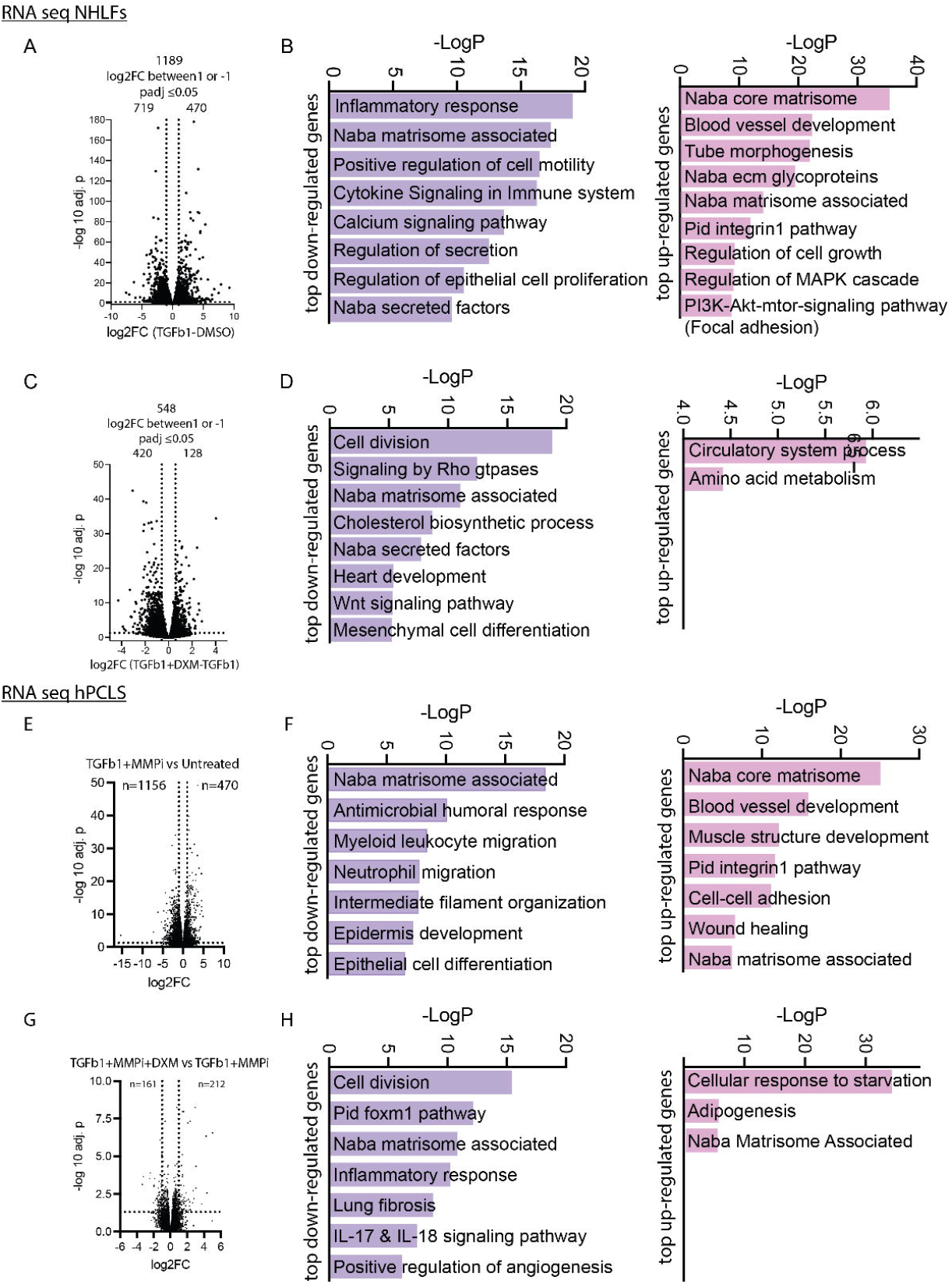
DXM elicits an anti-fibrotic transcriptional response in in-vitro and ex-vivo models of fibrogenesis. E, G, Volcano plot analysis of the differentially expressed genes (log2FC between 1 or −1, padj ≤ 0.05) in NHLFs treated with TGFb1 and TGFb1+DXM respectively. F, H, bar graph represents the enriched pathways elicited by top significantly up- and downregulated genes in respective treatment of NHLFs. E, C, volcano plot analysis of the differentially expressed genes (log2FC between 1 or −1, padj ≤ 0.05) in hPCLS treated either with TGFb1+MMPi or TGFb1+MMPi+DXM for 2 weeks of *ex-vivo* culture. B, D, the bar graphs represent pathways enriched in top up- and downregulated genes upon TGFb1+MMPi or TGFb1+MMPi+DXM treatment of hPCLS for 2 weeks of ex-vivo culture.

Co-treatment of NHLF cells with TGFb1 and DXM revealed a significant (log2FC between 1 or −1, padj ≤ 0.05) change in expression of 548 genes (Figure 6C) compared to cells treated with TGFb1 alone (Figure 6C-D). Of these 420 genes were down regulated and 128 were upregulated. All fibrosis associated pathways upregulated by TGFb1 treatment did not show significant upregulation after DXM treatment compared to control cells.

Additional pathways including Rho GTPases signalling, Wnt signalling, mesenchymal cell differentiation^1,9,42^ and cell division pathways were also downregulated by DXM compared to TGFb1 treatment (Figure 6D). Furthermore, co-treatment of NHLF with DXM and TGFb1, caused significant downregulation of pro-fibrotic collagens, *COL3A1, COL6A6, COL8A1, COL10A1* compared to control cells (Table-S2-Trancriptomics-NHLF). Data showed a significant downregulation of myofibroblast marker, ACTA2 (alpha smooth muscle actin, log2FC of −0.71) as well (Table-S2-Trancriptomics-NHLF). This observation prompted us to validate if DXM treatment has any effect on the myofibroblasts differentiation. Here, immunofluorescence analysis of NHLFs treated with TGFb1 stimulation, showed a robust cytoskeletal stress fiber signal of ACTA2 protein (Figure S4), consistent with the transcriptome analysis, indeed, this increase was significantly reduced upon co-treatment with TGFb1 and DXM (Figure S4). Altogether, these data suggest that DXM mediated anti-fibrotic effects may act via downregulation of pro-fibrotic ECM genes, blunting the onset of pro-fibrotic development program^43^ (WNT signalling) and preventing myofibroblasts differentiation of fibroblasts.

In addition, bulk mRNA sequencing analysis of fibrotic stimulated hPCLSs (TGFb1+MMPi)^21^ showed 470 upregulated and 1156 downregulated genes compared to control HPCLSs (Figure 6E-H) [Table-S3-Trancriptomics-hPCLS-TGFB-MMPI-Untreated]. Similarly, to our experiments with NHLF cells, TGFb1+MMPi stimulation of hPCLSs upregulates genes that elicit activity of pro-fibrotic pathways (Figure 6F) such as Naba Core Matrisome, Blood vessel development, wound healing, cell adhesion and Integrin pathway similar to that found in pulmonary fibrosis tissues^41^(Figure 6E-F). However, the significantly downregulated pathways overlapped only to those associated to Naba Matrisome. Matrisome genes that are crucial for pulmonary fibrosis progression and ECM remodeling such *COL1A1, COL1A2, COL3A1* and *COL5A1*^44^ (among others), *POSTN, THBS2*, *MMP2* and myofibroblast markers *ASPN, MYH11, POSTN, CNN1*^45^ showed upregulation (Table-S3-Trancriptomics-hPCLS-TGFB-MMPI-Untreated). Taken together, these data suggest that our TGFb1+MMPi treated hPCLS model system reflects the ECM remodeling and fibroblast to myofibroblast trans-differentiation phenotypes of the pulmonary fibrosis tissue.

Next, to understand the effect of DXM on TGFb1+MMPi induced pro-fibrotic signalling in hPCLSs, we analysed the mRNA transcriptome profile of hPCLS treated with TGFb1+MMPi and TGFb1+MMPi+DXM (Figure 6G-H). The analysis showed the DXM treatment to hPCLSs (co-treated with TGFb1+MMPi) differentially regulates 373 genes (Figure 6G) [161 downregulated and 212 upregulated]. Pathway enrichment of 161 downregulated genes included Naba matrisome associated pathway and down-regulation of Lung fibrosis pathway (Figure 6H). Upregulated genes where mainly associated to the response for amino acid deficiency. Gene targeted analysis of DXM treated hPCLSs showed significant downregulation of genes that have been shown to be upregulated in pulmonary fibrosis lung tissue such as *TGFBI, ADAMTS12, ELN and THBS1 (*Table-S5-Trancriptomics-hPCLS-TGFB-MMPI-DXM-TGFB-MMPI)^41,46^. Furthermore, pro-fibrotic genes such as *TGFBI, SERPINB2, IL24, HAS2, FGF1, S100A3, CNN1, AQP1, TAGLN, ELN, ADAMTS12, THBSI AND BGN* (Table-S5-Trancriptomics-hPCLS-TGFB-MMPI-DXM-TGFB-MMPI) are downregulated compared to control hPCLS (TGFb1+MMPi treated), providing further evidence to the potential anti-fibrotic activity of DXM.

## Discussion

Drug development, with subsequent approval for use in humans, is a challenging process because of the high costs involved across lengthy timelines^47^. This process is further made cumbersome due to lower success rates of lead molecules passing through various phases of clinical trials testing the respective toxicity and efficacy in patients. To this end, in general, re-purposing of FDA-approved drugs presents a potentially efficient solution to existing drug development problems^48^. This study identifies Dextromethorphan (DXM), an FDA-approved drug that is used in cough syrups, as a potential anti-pulmonary fibrosis drug, based on its efficacy in decreasing ECM-deposited COL1 (Figure 2A-G). Excessive, aberrant deposition of fibrillar collagen is the common pathological phenotype of lung fibrosis independent of its cause (genetic, environmental or idiopathic)^1^. Therefore, any compound inhibiting this process has the potential to ameliorate lung fibrosis. Furthermore, successful drug development against any disease involves efficacy testing in model systems close to human physiology and respective disease pathology. To this end, we have tested the potential anti-fibrotic efficacy of DXM in both *in-vitro* cultured NHLFs and *ex-vivo* cultured human PCLS, and both are established model systems of drug discovery against pulmonary fibrosis^16,49^.

Integrating Label-free SHG, secretome, and bulk mRNA sequencing analyses revealed that DXM elicits a multifactored anti-fibrotic response. It inhibits excess pro-fibrotic deposition of fibrillar collagens in ECM, reduces the abundance of pro-fibrotic collagens (such as COL1, COL3, COL5) and other core matrisome members (such as FBLN1, TNC, THBS2) in extracellular space. In addition, DXM also reduces the abundance of latent transforming growth factor binding proteins 1 and 2 (LTBP1, LTBP2) and TGFb1 pro-protein in extracellular space, which leads to inhibition of TGFb1 mediated pro-fibrotic signalling and the resulting activation of pathways (Figure 6) such as Naba core matrisome, integrin pathway, Wnt signalling and mesenchymal cell differentiation in both ex-vivo and in-vitro cultured models of pulmonary fibrosis.

In addition, Collagen Prolyl 4-Hydroxylases (P4Hs), stabilized by DXM as shown by the TPP analysis, is an established target for pulmonary fibrosis. Drugs that inhibit prolyl-hydroxylase enzymes, hence causing under-hydroxylation of collagens, also show a COL1 transport inhibition in ER, resulting in anti-fibrotic effects. Albeit, these inhibitors are used in high (300µM to ca. 7mM range) concentrations in-vitro as well in-vivo^50,51^ compared to the DXM concentration used in this study. As opposed to these inhibitors, DXM causes an increase in hydroxylation which results in transport inhibition in ER. Notably, COL1 accumulation in ER due to either lack of TANGO1^52^ or mutation in TANGO1^53^ and via pan-
hydroxylase inhibitor dimethyloxalylglycine (DMOG)^54^ causes downregulation of COL1 at the mRNA level. Furthermore, lack of TANGO1 in mice with CCL4-induced liver fibrosis blunts upregulation of pro-fibrotic markers such as ACTA2, TIMP1, TGFb and PDGFRß^55^. In case of TGFb1-treated hepatic stellate cells, co-treatment with DMOG reduces COL1 expression as well and blunts the TGFb1-induced activation of pro-fibrotic cells (as shown by decreased *ACTA2* expression)^54^. These studies suggest that, indeed a block of collagen transport in ER via DXM in addition to LTBP1 and LTBP2 adds to the anti-fibrotic potential of DXM.

Furthermore, this mechanism of action is different compared to existing anti-fibrotic drugs i.e., Nintedanib and Pirfenidone. Nintedanib causes an inhibition PDGF, VEGF and FGF signalling^56^, Pirfenidone on the other hand causes an anti-fibrotic effect by inhibiting TNFalpha translation and GLI transcription factor activity^57,58^. This raises the possibility of combinatorial treatment of pulmonary fibrosis patients using DXM together with Nintedanib or Pirfenidone, which could potentially improve the efficacy of current treatment regimen. Additionally, previous studies have shown DXM to be effective in UUO mouse model of kidney fibrosis^59^. These studies together with our detailed mechanism of action advocates for the use of DXM as an anti-fibrotic drug in other fibrotic organs such as liver, kidney, heart and skin as well.

The molecular target that causes DXM-induced collagen, LTBP1, LTBP2 transport inhibition in the ER still remains to be investigated. SIGMAR1 is one of the main known targets of DXM^20^ and localizes at the endoplasmic reticulum^20^. SIGMAR1 antagonists (S1RA^60^ or BD1047^61^) or agonist (PRE-084^62^) could not induce an ER inhibition of COL1 transport (Khan et al., unpublished results) in NHLF cells, suggesting a target of DXM other than SIGMAR1 may be responsible for the transport inhibition of COL1 upon DXM treatment. To this end, target engagement analysis using thermal proteome profiling revealed that DXM causes a stabilization of ER resident proteins important for COL1 folding (Figure 5B, black dashed rectangle). Notably, these proteins include prolyl-hydroxylases (P3H2, P3H3, P3H4, P4HA1, P4HA2), prolyl-hydroxylase complex associated protein CRTAP and lysyl hydroxylase PLOD1 and PLOD2 (Figure 5B). Consistent with the stabilization of hydroxylase enzymes, hydroxylation level of collagen (COL1, COL3, COL4, COL5, COL6, COL7, COL12), LTBP1 and LTBP2 peptides detected in cell lysate of TGFb1+DXM treated NHLFs were significantly increased (Figure 5C) compared to TGFb1 treatment.

These data suggest that DXM mediated hyper hydroxylation of collagen and like cargoes is the likely cause for their transport inhibition. In support of this hypothesis, previous studies have shown that hydroxylation represents a critical step during folding and thus transport of secretory cargoes such as collagens^40^.

Integrating mass spectrometry analysis of NHLFs supernatants and cell lysates using thermal proteome profiling and hydroxylation analysis revealed that DXM-induced trafficking inhibition occurs preferably in secreted cargoes that are hyper hydroxylated (Figure 3H, 5B-C). Most of these proteins include core matrisome members (COL1, COL3, COL4, COL5, COL6, COL7, COL12, LTBP1 and LTBP2). These data suggest the existence of a cell secretion mechanism that might regulate the transport of a subset proteins involved in extracellular remodelling. Further experiments would be needed to reveal the exact molecular target to which DXM binds to regulate these proteins. Once this target is known, the resulting knowledge could shed light on mechanisms going beyond fibrosis to other diseases and pathways such as tumour formation, wound repair, tissue morphogenesis, differentiation and homeostasis^63^.

Furthermore, DXM-induced targeted inhibition of collagen depostion in extracellular matrix could be used as an excellent cell biological tool to control the composition of extracellular matrix, that could culminate in forming extracellular matrix of defined mechanical properties as well. This property of DXM will find ample applications in the field of matrix biology in general. Since, DXM mediated collagen transport block in the ER is reversible and fast (Figure 3G, Figure 4C-D), this provides a unique opportunity of studying the kinetics of endogenous collagens and like cargoes. In conjunction with high throughput screening technology, results will reveal novel regulators of transport of endogenous collagens, LTBP1, LTBP2 and other like cargoes.

To our knowledge, DXM is the first drug that shows trafficking based inhibition of TGFb1 activity regulating proteins (LTBP1, LTBP2 and TGFb1 proprotein). This property will help target, understanding paracrine and autocrine TGFb1 signalling in diverse physiological and pathological processes. Furthermore, pulldown assays together with mass spectrometry analysis of these cargoes before and after DXM treatment, could be used to reveal novel post-translational modifications of the respective secreted proteins that might control disease pathology.

In summary, reversible trafficking inhibition of collagens, LTBP1, LTBP2, TGFb1 pro-proteins, down-regulation of anti-fibrotic pathways by DXM in different models of fibrogenesis provides compelling evidence that DXM could prove to be a successful ECM targeted therapy for treating pulmonary fibrosis or fibrosis in general in the future.

## Materials and Methods

### Ethical use of human biomaterial

Tumour-free lung tissue (tissue resections of lung cancer patients) was obtained from Thoraxklinik-Heidelberg (anonymised patient identities). In charge pathologist designated the tissue as tumour-free. Patient consent and use of tissue were obtained as per the research ethics committee (Medical Faculty of University Heidelberg and EMBL BIAC committee) under approval reference number S-270/2001 and BIAC 2021-003. Primary normal (CC-2512) and IPF (CC-7231) human lung fibroblasts were purchased from Lonza. Human biological samples were sourced ethically and their research use was in accordance with the terms of informed consent under the institutional review board/ethics committee-approved protocol. hTERT-immortalized BJ-5ta cells (#CRL-4001) were purchased from ATCC. Pericytes, PDGFRb+ mesenchymal cells were a kind gift from Prof. Rafael Kramann, RWTH Aachen.

### FDA-approved drug library screening assay

The library of 762 drugs (Table-S8-DrugLibraryList) was used to screen for molecules that could be used for treating pulmonary fibrosis. The drugs were dissolved in DMSO. Using cell seeder (ThermoFisher, Multidrop Combi #5840300) primary normal human lung fibroblasts were plated at a density of 6000 cells per well of a 96 well glass bottom plate (Zellkontakt #5241). The cells were seeded in Lonza FBM-FCS fibroblast media (#CC-3131, #CC-4126). 18 hours post seeding, the media was replaced with DMEM (Gibco #31885-023) with a mixture of Ficol 70 (Sigma #F2878) and 400 (Sigma #F4375). Additionally, all the wells were treated with TGFb1 (R&D systems, #240-B-010, 5ng/ml). Drugs were used at a concentration of 10µM, except for positive controls (Nintedanib, 5µM and Pirfenidone, 0.5mM). 48 hours post-stimulation (±TGFb1 and Drug), the cells were directly fixed in 3% PFA (10 minutes). Washed with PBS (3X). Next, the cells were incubated with a primary antibody against COL1 (0.2µg/ well, Abcam #ab34710) in PBS for 1 hour and 30 minutes. Subsequently, cells were washed (using Tecan Hydrospeed #30054550) with PBS (3 times) and incubated with a secondary antibody (1:500, Molecular Probes AF488 #A11008) and Hoechst33342 (1:2000, ThermoFisher #H21492). The 96 well plates were imaged using a 20X air objective in a widefield microscope set up (Molecular Devices IXM, CFI P-Apo 20X Lambda/ 0.75/ 1,00). Each of the plate was barcoded, the images were analysed using Cell-profiler software, robust z-scores were calculated using shinyHTM^64^. Robust z-score quantifying the relative effect of each stimulation (TGFb1 or TGFb1+Drug) was calculated in different biological replicate and according to the following formula: Robust z-score score = xi – X/ MAD, where xi is the average extracellular matrix COL1 signal per nucleus in the respective image of the well, and X is determined as the median extracellular matrix COL1 signal per nucleus of the whole plate. MAD is the median absolute deviation in the whole pate. Negative robust z-scores represent decrease in deposited COL1 in extracellular matrix, and positive robust z-scores represent increase in deposited COL1 in extracellular matrix. To select top anti- or pro-collagen deposition hits, drugs were also filtered for the average number of nuclei per image. Only drugs that maintained nuclei number ≥ 120 were selected. This was done to avoid effects on collagen deposition due to potential cell death that the respective drug may have caused.

### Immunofluorescence staining and confocal microscopy

Cells were fixed in 3% PFA (10 minutes, invariably). Washed with PBS (3X). Permeabilized with 0.1% Triton X-100 (in PBS for 10 minutes). Washed with PBS (3X). Incubated with respective primary and secondary antibodies (1 hour 30 minutes each), with PBS washed in between incubations. Different antibodies used are; COL1 (Rockland# 600-401-103-0.5; Merck# C2456), TANGO1 (Invitrogen# PA5-63209), HSP47 (Enzo# ADI-SPA-470-D), GM130 (BD Biosciences#610822), Calnexin (Enzo#ADI-SPA-860F), ERGIC53 (Enzo#ALX-804-602-C100), PDI (Cell Signalling#3501), P3H1 (Novus#H00064175-B01P), ASMA (Abcam# 76549). Images were acquired using NIKON A1 and Zeiss LSM 780 confocal microscopes. *X meaning, how many times a procedure was repeated. Drugs and Chemicals tested: Dextromethorphan HBr (Sigma#D9684),

### Transmission Electron Microscopy of in-vitro cultured NHLFs

Cells were fixed with 2% glutaraldehyde in 0.1 M sodium cacodylate buffer pH 7.4 at room temperature for 2 hours. Following this, cells were washed (6 × 10 min) with 0.1 M sodium cacodylate buffer and post-fixed in reduced osmium tetroxide (1% OsO4/ 1.5% K3Fe (III)(CN) in sodium cacodylate buffer pH 7.4) for 2 hours at 4°C in the dark. Cells were then washed again (6 × 10 min) with distilled water and gradually dehydrated with increasing concentrations of ethanol; 50%, 70%, 90% and 4×100% at RT and were gradually infused with an Epon 812-ethanol mixture; 25%, 50%, 75%, 2 x pure Epon. After embedding in pure Epon 812 resin samples were polymerized for 48 hours at 65°C. Ultrathin sections of 70 nm were cut using a diamond knife (Ultra 45° Diatome) on a Leica Ultracut UCT7, placed on 50 mesh copper grids with carbon coated formvar support film and stained with uranyl acetate and Waltons lead citrate. High resolution EM imaging was done on a Jeol 2100Flash TEM at 120 KeV. All electron microscopy imaging was completed at the EM Core Facility, EMBL-Heidelberg.

### hPCLS ex-vivo culture

Tumour free resected human lung tissue was obtained from Thoraxklinik Heidelberg, Germany. Lung tissue was inflated with 3% agarose and prepared as described previously^21^. Post inflation, cylindrical cores of lung tissue were prepared using Krumdieck tissue coring press tool (#MD5000). The tissue cylinders were cut in to 350µm thick and 5 mm (dimeter) precision-cut tissue slices using a Leica vibratome VT 1200 S.

### Label-free SHG imaging

To visualize SHG signal from ECM-deposited by NHLFs, cells were cultured for 8 days in glass bottom 24 well plates (15,000 cell per well on day 0), with cell culture media replenishment after every 48 hours. The cells were chemically fixed in 3% PFA and washed with PBS. SHG imaging was performed as described previously^21^ using a Zeiss LSM 780-NLO microscope.

### Mass spectrometric analysis of in-vitro cultured NHLFs

LC-MS/MS analysis of lysates. 20 µg of lysates were subjected to an in-solution tryptic digest using a modified version of the Single-Pot Solid-Phase-enhanced Sample Preparation (SP3) protocol^65,66^.

In total three biological replicates were prepared including mock (DMSO), TGFB1, Dextromethorphan as well as and TGFB1/Dextromethorphan treated cells (n=3). Lysates were added to Sera-Mag Beads (Thermo Scientific, #4515-2105-050250, #6515-2105-050250) in 10 µl 15% formic acid and 30 µl of ethanol. Binding of proteins was achieved by shaking for 15 min at room temperature. SDS was removed by 4 subsequent washes with 200 µl of 70% ethanol. Proteins were digested overnight at room temperature with 0.4 µg of sequencing grade modified trypsin (Promega, #V5111) in 40 µl Hepes/NaOH, pH 8.4 in the presence of 1.25 mM TCEP and 5 mM chloroacetamide (Sigma-Aldrich, #C0267). Beads were separated, washed with 10 µl of an aqueous solution of 2% DMSO and the combined eluates were dried down. Peptides were reconstituted in 10 µl of H2O and reacted for 1 h at room temperature with 80 µg of TMT6plex (Thermo Scientific, #90066) label reagent dissolved in 4 µl of acetonitrile. Excess TMT reagent was quenched by the addition of 4 µl of an aqueous 5% hydroxylamine solution (Sigma, #438227). Peptides were reconstituted in 0.1 % formic acid and mixed to achieve a 1:1 ratio across all TMT-channels. Thus, mixed peptides were purified by a reverse phase clean-up step (OASIS HLB 96-well µElution Plate, Waters #186001828BA). Peptides were subjected to an off-line fractionation under high pH conditions. The resulting 12 fractions were then analyzed by LC-MS/MS on an Orbitrap Fusion Lumos mass spectrometer (Thermo Scentific). To this end, peptides were separated using an Ultimate 3000 nano RSLC system (Dionex) equipped with a trapping cartridge (Precolumn C18 PepMap100, 5 mm, 300 μm i.d., 5 μm, 100 Å) and an analytical column (Acclaim PepMap 100. 75 × 50 cm C18, 3 mm, 100 Å) connected to a nanospray-Flex ion source. The peptides were loaded onto the trap column at 30 µl per min using solvent A (0.1% formic acid) and eluted using a gradient from 2 to 38% Solvent B (0.1% formic acid in acetonitrile) over 90 min at 0.3 µl per min (all solvents were of LC-MS grade). The Orbitrap Fusion Lumos was operated in positive ion mode with a spray voltage of 2.4 kV and capillary temperature of 275 °C. Full scan MS spectra with a mass range of 375–1500 m/z were acquired in profile mode using a resolution of 60,000 (maximum fill time of 50 ms and a RF lens setting of 30%. Fragmentation was triggered for 3 s cycle time for peptide like features with charge states of 2–7 on the MS scan (data-dependent acquisition). Precursors were isolated using the quadrupole with a window of 0.7 m/z and fragmented with a normalized collision energy of 36%. Fragment mass spectra were acquired in profile mode and a resolution of 15,000. Maximum fill time was set to 54 ms. The dynamic exclusion was set to 60 s. Acquired data were analyzed using IsobarQuant^67^ and Mascot V2.4 (Matrix Science) using a reverse UniProt FASTA Homo sapiens database (UP000005640) including common contaminants. The following modifications were taken into account: Carbamidomethyl (C, fixed), TMT10plex (K, fixed), Acetyl (N-term, variable), Oxidation (M, variable) and TMT10plex (N-term, variable). The mass error tolerance for full scan MS spectra was set to 10 ppm and for MS/MS spectra to 0.02 Da. A maximum of 2 missed cleavages were allowed. A minimum of 2 unique peptides with a peptide length of at least seven amino acids and a false discovery rate below 0.01 were required on the peptide and protein level^68^. The raw output files of IsobarQuant (protein.txt – files) were processed using the R programming language (ISBN 3-900051-07-0). Only proteins that were quantified with at least two unique peptides were considered for the analysis. Moreover, only proteins which were identified in two out of three mass spectrometric runs were kept. 5899 proteins passed the quality control filters. Raw TMT reporter ion intensities (‘signal_sum’ columns) were first cleaned for batch effects using limma^69^ and further normalized using vsn (variance stabilization normalization^70^. Missing values were imputed with ‘knn’ method using the Msnbase package^71^. Proteins were tested for differential expression using the limma package. The replicate information was added as a factor in the design matrix given as an argument to the ‘lmFit’ function of limma. Also, imputed values were given a weight of 0.05 in the ‘lmFit’ function. A protein was annotated as a hit with a false discovery rate (fdr) smaller 0.05 and a fold-change of at least 100 % and as a candidate with an fdr below 0.02 and a fold-change of at least 50 %.

### Thermal Proteomic Profiling

Thermal proteomic profiling has been performed according to Becher et al^72^. Briefly, 15-cm dishes of NHLF cells were cultured for 1 h in medium supplemented with fetal calf serum in the absence or presence of DXM (0, 1, 5, 10µM final concentration). All following steps were carried out in the presence of vehicle (DMSO) or DXM (1, 5, 10µM final concentration). Cells were washed twice with PBS and trypsin was added. Cells were harvested in DMEM supplemented with fetal calf serum. Cells were washed twice with PBS and counted. 14×10^6 cells were transferred to a 15ml tube, spun down (1.000 rpm, 2 min, room temperature), the supernatant was removed and the cells were resuspended in 1.4 ml PBS supplemented with the vehicle or the drug to achieve a suspension of 1×10^6 cells/100 µl. 1×10^6 cells were transferred to a PCR plate and cells were spun down. 80 µl of the supernatant were removed. The PCR plate was sealed with aluminum foil (Mircroplate Foil, 96 well, GE Healthcare, # 28-9758-16) and a temperature gradient of 37°C to 67°C was applied for 3 min using a PCR machine. The plate was removed and transferred to room temperature. After 3 min, the plate was transferred on ice and samples were allowed to cool down for 3 min. 30 µl of lysis buffer were added and samples were resuspended by pipetting. Samples were lysed for 1 h at 4°C while shaking at 500 rpm. A filter plate (MultiScreen, 96-well Plate, 0.45 µm, REF #MSHVN4550) was equilibrated by the addition of 50 µl plate wash solution to each well followed by centrifugation at 2.000 rpm for 3 min at 4°C. Lysates were subjected to centrifugation at 2.000 rpm for 3 min at 4°C step to remove aggregates. 45 µl of the supernatants were transferred to the filter plate. Aggregates were removed by centrifugation at 500 g for 10 min at 4°C. 25 µl of the filtrate was transferred to a new plate and 25 µl of a 2x sample buffer were added. A BCA protein determination on remaining samples was performed. 37°C samples were adjusted to 10 mg/ml by the addition of 1x sample buffer. The same volume of 1x sample buffer has been added to all other samples of the respective condition.

### Sample preparation for the mass spectrometric analysis

10 µg of protein were subjected to an in-solution tryptic digest using a modified version of the Single-Pot Solid-Phase-enhanced Sample Preparation (SP3) protocol^65,66^. 1% SDS-containing lysates were added to Sera-Mag Beads (Thermo Scientific, #4515-2105-050250, 6515-2105-050250) in 10 µl 15% formic acid and 30 µl of ethanol. Binding of proteins was achieved by shaking for 15 min at room temperature. SDS was removed by 4 subsequent washes with 200 µl of 70% ethanol. Proteins were digested overnight at room temperature with 0.4 µg of sequencing grade modified trypsin (Promega, #V5111) in 40 µl Hepes/NaOH, pH 8.4 in the presence of 1.25 mM TCEP and 5 mM chloroacetamide (Sigma-Aldrich, #C0267). Beads were separated, washed with 10 µl of an aqueous solution of 2% DMSO and the combined eluates were dried down. Peptides were reconstituted in 10 µl of H2O and reacted for 1 h at room temperature with TMT10 (Thermo Scientific, #90110) or TMTpro 18-plex (Thermo Scientific, #A44522) labelling reagent. To this end, 80 µg of TMT10 or 100 µg of TMTpro 18-plex label reagent were dissolved in 4 µl of acetonitrile and added to the peptides. Excess TMT reagent was quenched by the addition of 4 µl of an aqueous 5% hydroxylamine solution (Sigma, 438227). Peptides were reconstituted in 0.1 % formic acid and equal volumes were mixed. Mixed peptides were purified by a reverse phase clean-up step (OASIS HLB 96-well µElution Plate, Waters #186001828BA). Peptides were subjected to an off-line fractionation under high pH conditions^65^. TMT10-labelled TPP samples as well as samples of the extracellular matrix 6 fractions were analyzed by LC-MS/MS using a 90 min gradient. For the corresponding full proteome 12 fractions were analyzed by LC-MS/MS applying a 120 min gradient.

### Analysis of TMT10-labelled TPP samples

Peptides were separated using an Ultimate 3000 nano RSLC system (Dionex) equipped with a trapping cartridge (Precolumn C18 PepMap100, 5 mm, 300 μm i.d., 5 μm, 100 Å) and an analytical column (Acclaim PepMap 100. 75 × 50 cm C18, 3 mm, 100 Å) connected to a nanospray-Flex ion source. The peptides were loaded onto the trap column at 30 µl per min using solvent A (0.1% formic acid) and eluted using a gradient from 2 to 38% Solvent B (0.1% formic acid in acetonitrile) over 82 min and then to 80% at 0.3 µl per min (all solvents were of LC-MS grade). The Orbitrap Fusion Lumos was operated in positive ion mode with a spray voltage of 2.4 kV and capillary temperature of 275 °C. Full scan MS spectra with a mass range of 375–1500 m/z were acquired in profile mode using a resolution of 120,000 with a maximum injection time of 50 ms, AGC operated in standard mode and a RF lens setting of 30%.

Fragmentation was triggered for 3 s cycle time for peptide like features with charge states of 2–7 on the MS scan (data-dependent acquisition). Precursors were isolated using the quadrupole with a window of 0.7 m/z and fragmented with a HCD collision energy of 36%. Fragment mass spectra were acquired in profile mode and a resolution of 30,000. Maximum injection time was set to 94 ms or a normalized AGC target of 200%. The dynamic exclusion was set to 60 s.

### Data analysis thermal proteomic profiling

Data were analyzed using IsobarQuant^67^ and Mascot V2.4 (Matrix Science) using a reverse UniProt FASTA Homo sapiens database (UP000005640 with 92607 entries, May 14^th^ 2016) including common contaminants. The following modifications were taken into account: Carbamidomethyl (C, fixed), TMT10plex (K, fixed), Acetyl (N-term, variable), Oxidation (M, variable) and TMT10plex (N-term, variable). The mass error tolerance for full scan MS spectra was set to 10 ppm and for MS/MS spectra to 0.02 Da. A maximum of 2 missed cleavages were allowed. A minimum of 2 unique peptides with a peptide length of at least seven amino acids and a false discovery rate below 0.01 were required on the peptide and protein level^68^.

### Analysis of TMTpro 18-plex labelled samples of the Secretome and the corresponding Cell lysate proteome

NHLFs were cultured in DMEM (FCS free) for 48 hrs. Cells were treated with TGFb1 or TGFb1+DXM. Subsequently, media supernatants were harvested, spun down at 1000 rpm for 5 minutes to remove dead cells and debri. The media supernatant was precipitated using TCA precipitation protocol^73^. The protein pellet of each condition was resuspended in 1% SDS in 100mM HEPES-NaOH. The respective cell lysates were prepared similarly as described under “Mass spectrometric analysis of in-vitro cultured NHLFs”. Furthermore, in solutiuon digestion steps were also performed similarly as described in “Mass spectrometric analysis of in-vitro cultured NHLFs”. Next, Peptides were separated using an Ultimate 3000 nano RSLC system (Dionex) equipped with a trapping cartridge (Precolumn C18 PepMap100, 5 mm, 300 μm i.d., 5 μm, 100 Å) and an analytical column (Acclaim PepMap 100. 75 × 50 cm C18, 3 mm, 100 Å) connected to a nanospray-Flex ion source. The peptides were loaded onto the trap column at 30 µl per min using solvent A (0.1% formic acid) and eluted using a gradient from 2 to 80% Solvent B (0.1% formic acid in acetonitrile) over 90 (extracellular matrix) or 120 min (full proteome of extracellular matrix) at 0.3 µl per min (all solvents were of LC-MS grade). The Orbitrap Fusion Lumos was operated in positive ion mode with a spray voltage of 2.2 kV and capillary temperature of 275 °C. Full scan MS spectra with a mass range of 375–1500 m/z were acquired in profile mode using a resolution of 120,000 with a maximum injection time of 50 ms, AGC operated in standard mode and a RF lens setting of 30%.

Fragmentation was triggered for 3 s cycle time for peptide like features with charge states of 2–7 on the MS scan (data-dependent acquisition). Precursors were isolated using the quadrupole with a window of 0.7 m/z and fragmented with a normalized collision energy of 34%. Fragment mass spectra were acquired in profile mode and a resolution of 30,000. Maximum injection time was set to 94 ms and AGC target to custom. The dynamic exclusion was set to 60 s.

### Data analysis of secretome and the corresponding total proteome

Raw files were analyzed with FragPipe v19.2 (MS-Fragger v3.7, IonQuant v1.8.10 and philosopher v4.8.1) using a UniProt FASTA database (UP000005640, 20594 entries, October 26th 2022). Standard setting of FragPipe’s ‘TMT16’ workflow were used. ‘Label type’ was changed to ‘TMT-18’ under the ‘Quant (Isobaric)’ tab. Under the ‘Validation’ tab, ‘Run PSM Validation’ was set to ‘Run PeptideProphet’ and ‘Run PTMProphet’ was activated (Cmd line opts: --keepold --static --em 1 --nions b --mods M:15.994900,KP:15.9949 --minprob 0.5). The following modifications were taken into account: Carbamidomethyl (C, fixed), TMT18plex (K, fixed), Acetyl (N-term, variable), Oxidation (M, variable) and TMT10plex (N-term, variable), Hydroxylation (P and K, variable).

### mRNA sequencing of in-vitro cultured NHLFs

Total RNA was extracted using RNAeasy kit (Qiagen). mRNA-seq libraries were generated using NEB Ultra II stranded mRNA chemistry (NEBNext® Ultra™ II DNA Library Prep Kit for Illumina®), as per the manufacturers protocol with the following settings: 500 ng total RNA input material, a fragmentation time of 8 minutes. The adaptor dilution was kept at 1:20. The samples were individually barcoded with NEBs dual indices and amplified for 8 PCR cycles. The library prep was performed on the i7 Beckman Liquid Handling System. The libraries were pooled and sequenced on the NextSeq 500, using the High Output 75 cycles kit.

### mRNA sequencing of ex vivo cultured hPCLS

The slices were stored in the freezer at −80°C until RNA isolation. The total RNA was isolated from each slice separately using an adapted protocol from RNeasy Mini Kit (RNeasy Mini Kit #74104, Qiagen). Each slice was placed into a 2ml low binding tube (DNA LoBind® #0030108078, #0030108051) containing 150ul Trizol (Invitrogen™ #15596026) and a 5mm stainless steel bead (#69989, Qiagen). The tubes were then fitted into a tissue homogenizer (TissueLyser II, # 85300, Qiagen) and homogenized through 3 rounds of 1min at 27 Hz and room temperature. The homogenate was transferred into a new 1.5ml low binding tube and 100ul of 100% Ethanol was added and mixed by pipetting. The lysate was then transferred, including precipitate if any, to a RNeasy Mini spin column in a 2ml collection tube and centrifuged (all centrifugation steps in this protocol were at 8000 x g and for 30 seconds, unless indicated otherwise and after each centrifugation flow through was discarded, except the last elution step). This was followed by adding 700ul of Buffer RW1 to the column and centrifuged. 500ul of Buffer RPE was added to the column and centrifuged – this step was repeated twice. Additional centrifugation was done for 2min to dry the membrane. In order to degrade the DNA, a DNase I solution was prepared by mixing 10ul of DNase I stock solution with 70ul RDD buffer and this mix was added directly to the column membrane. The incubation was carried out for 15min at room temperature. After the incubation, 350ul of buffer RW1 was added to the RNeasy column and centrifuged. 500ul of Buffer RPE was added to the column and centrifuged. The RNeasy spin column was placed in a new 1.5ml collection tube and 20ul of RNase free water was added directly to the membrane and incubated for 1min. Then the RNeasy spin column was centrifuged for 1min to elute the RNA. The quality of the RNA was evaluated on Qbit and Bioanalyzer using Agilent RNA Pico kit. **Library preparation,** the extracted RNA was quantified using the Invitrogen Qubit 4, with the RNA HS Assay as per the manufacturer’s protocol. For the measurement 1ul of sample in 199ul of Qubit working solution was used. Based on these results the integrity of the RNA was analysed using the Agilent Bioanalyzer with the RNA Pico Assay kit as per the manufacturer’s protocol. Based on the quality assessment, the samples were then standardized to 1 ng of RNA in a volume of 2.4 ul. For library preparation a modified version of the *Picelli, S. et al. (2014)*^74^ *and Henning et al. (2018)*^75^ was used. The protocol was started at step 5 of the paper Picelli et al. using 2.4ul of standardized RNA sample instead of 2ul of lysis-buffer. For reverse transcription SuperScript IV was used, to improve the reverse transcription efficiency. To compensate the higher starting volume, the H2O volume of the RT-Mix (step 10) was increased to 1.1ul. Using the SuperScript IV the incubation time in Step 11 was adjusted accordingly (98°C for 15 mins, 80°C for 10 min, cool down to 10°C). For Pre-amplification (Step 14), 15 cycles were used. After checking the quality of the cDNA in step 27, the cDNA was diluted to 0.2ng/ul. The diluted cDNA was processed following Henning et al. 2018 Supplemental Methods, Tn5(R27S), E54K, L372P loading and tagmentation-based NGS library preparation, using the SDS option for inactivation and the KAPA HiFi HotStart polymerase with 12 PCR cycles. The libraries were quantified using the Qubit HS DNA assay as per the manufacturer’s protocol. For the measurement 1ul of sample in 199ul of Qubit working solution was used. The quality and molarity of the libraries was assessed using Agilent Bioanalyzer with the DNA HS Assay kit as per the manufacturer’s protocol. The assessed molarity was used to equimolar combine the individual libraries into one pool for sequencing. The pool has been cleaned up using SPRI beads with a ratio of 0.7x as per the manufacturer’s protocol (Left Side Size Selection). After the clean-up the pool was quantified using the Qubit HS DNA assay and the size was assessed using the Agilent Bioanalyzer. For Sequencing it was loaded and sequenced on an Illumina NextSeq 2000 platform (Illumina, San Diego, CA, USA) using a P3 50 cycle kit and a read-length of 80 bases.

#### Sequencing Analysis

Reads were mapped to the human reference genome (GRCh38) using STAR (version 2.7.9a) aligner^76^ with default parameters. Number of counts per genes were produced during alignments (--quantMode GeneCounts) with the gene annotation version GRCh38.93. Differentially expressed genes were identified using DESeq2 (version 1.36.0).

## Supporting information

Table-S1-Proteomics-NHLF

Table-S2-Trancriptomics-NHLF

Table-S5-Trancriptomics-hPCLS-TGFB-MMPI-DXM-TGFB-MMPI

Table-S8-DrugLibraryList

Table-S9-DrugRobustZScores

Table-S11-Secretome-TGFb1+DXM-TGFb1-NHLF

Table-S12-OH-TGFb1+DXM-TGFb1-NHLF

Table-S3-Trancriptomics-hPCLS-TGFB-MMPI-Untreated

## Acknowledgments

We sincerely thank Ms. Christa Stolp for providing helpful assistance in ensuring a smooth availability and processing of human lung tissue. We sincerely thank Claudia A. Staab-Weijnitz (CPC Munich) for her advice on improving mass spectrometric coverage of collagens in our data.

## Figure Legends

**Figure S1.** A, representative images of ECM deposited COL1 by *in-vitro* cultured NHLFs in DMEM cell culture media, DMEM cell culture media together with Ficoll mixture (MMC=Macromolecular crowding) and DMEM cell culture media together with MMC and TGFb1 stimulation (MMC+TGFb1). B, quantification of ECM deposited COL1 in the respective conditions of NHLFs in vitro culture. C, representative immunofluorescence staining images of VSVG-YFP protein and VSVG protein present at the membrane of NHLFs. VSVG-YFP protein was expressed in NLHFs (green signal), this signal can be visualized both inside the cell and on the membrane. While VSVG membrane (red) signal, is obtained by staining non-permeabilized NHLFs with an antibody against VSVG protein. The cells were cultured over night at 40°C for transport block from ER and subsequently on 32°C for release of respective cargo from ER. D, quantitative analysis of VSVG transport ratio (signal on the membrane vs total signal).

**Figure S2.** A. immunofluorescence images of CCBE1 using confocal microscopy from NHLFs cultured in vitro upon stimulation with TGFb1 and TGFb1+DXM. B, Representative images are a single optical plane belonging to the respective z-stack of images. Immunofluorescence images of PDI and COL1 acquired using confocal microscopy from NHLFs cultured in vitro upon stimulation with TGFb1 and TGFb1+DXM. C, immunofluorescence images of SEC31 (ER exit site marker) and COL1 acquired using confocal microscopy from NHLFs cultured in vitro upon stimulation with TGFb1 and TGFb1+DXM. Scale bar = 50µm. n=2.

**Figure S3.** Representative images are a single optical plane belonging to the respective z-stack of images. A, Immunofluorescence images of different GM130 (Golgi complex marker, false colored as Green) and COL1 (false colored as red) acquired using confocal microscopy from NHLFs cultured in vitro upon stimulation with TGFb1 and TGFb1+DXM. B, Immunofluorescence images of Calnexin (ER marker, false colored as Green) and COL1 (false colored as red) acquired using confocal microscopy from NHLFs cultured in vitro upon stimulation with TGFb1 and TGFb1+DXM. C, Immunofluorescence images of ERGIC53 (Golgi-ER intermediate compartment marker, false colored as Green) and COL1 (false colored as red) acquired using confocal microscopy from NHLFs cultured in vitro upon stimulation with TGFb1 and TGFb1+DXM. Scale bar = 50µm. n=2

**Figure S4:** A, Representative maximum projection images of immunofluorescence staining against alpha smooth muscle actin (ASMA) in human embryonic primary fibroblasts (WI-38). The cells were treated with DMSO, TGFb1 and TGFb1+DXM. Scale bar =50µm. B, quantification of ASMA signal present on stress fibres normalized to the nuclei number per image. One-way ANOVA was used for testing statistical significance. p-value ≤0.001****. N=3.

