## Supplementary figures and images for "Dextromethorphan inhibits collagen transport in the endoplasmic reticulum eliciting an anti-fibrotic response in *ex-vivo* and *in vitro* models of pulmonary fibrosis"

### Figure S1

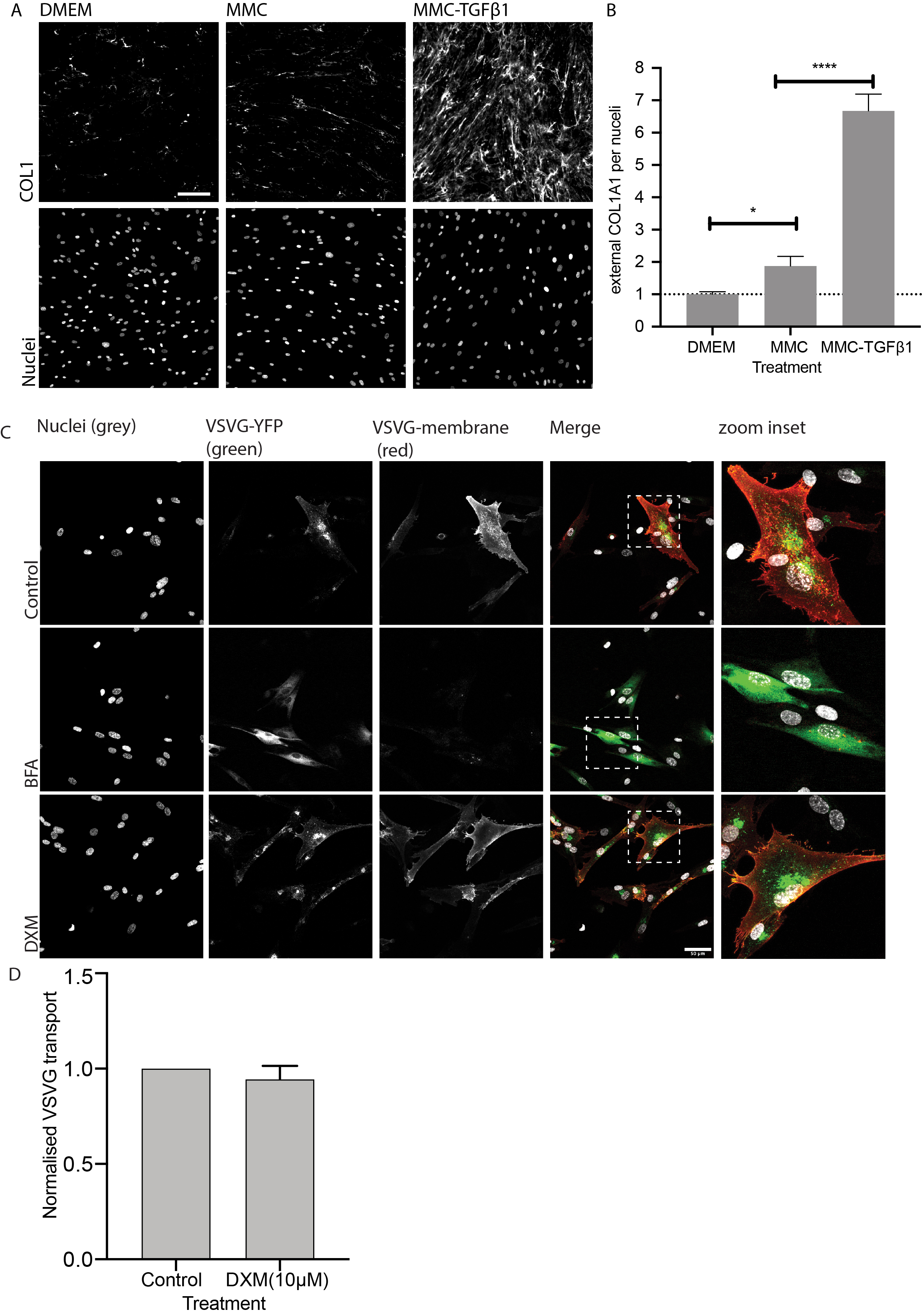

### Figure S2

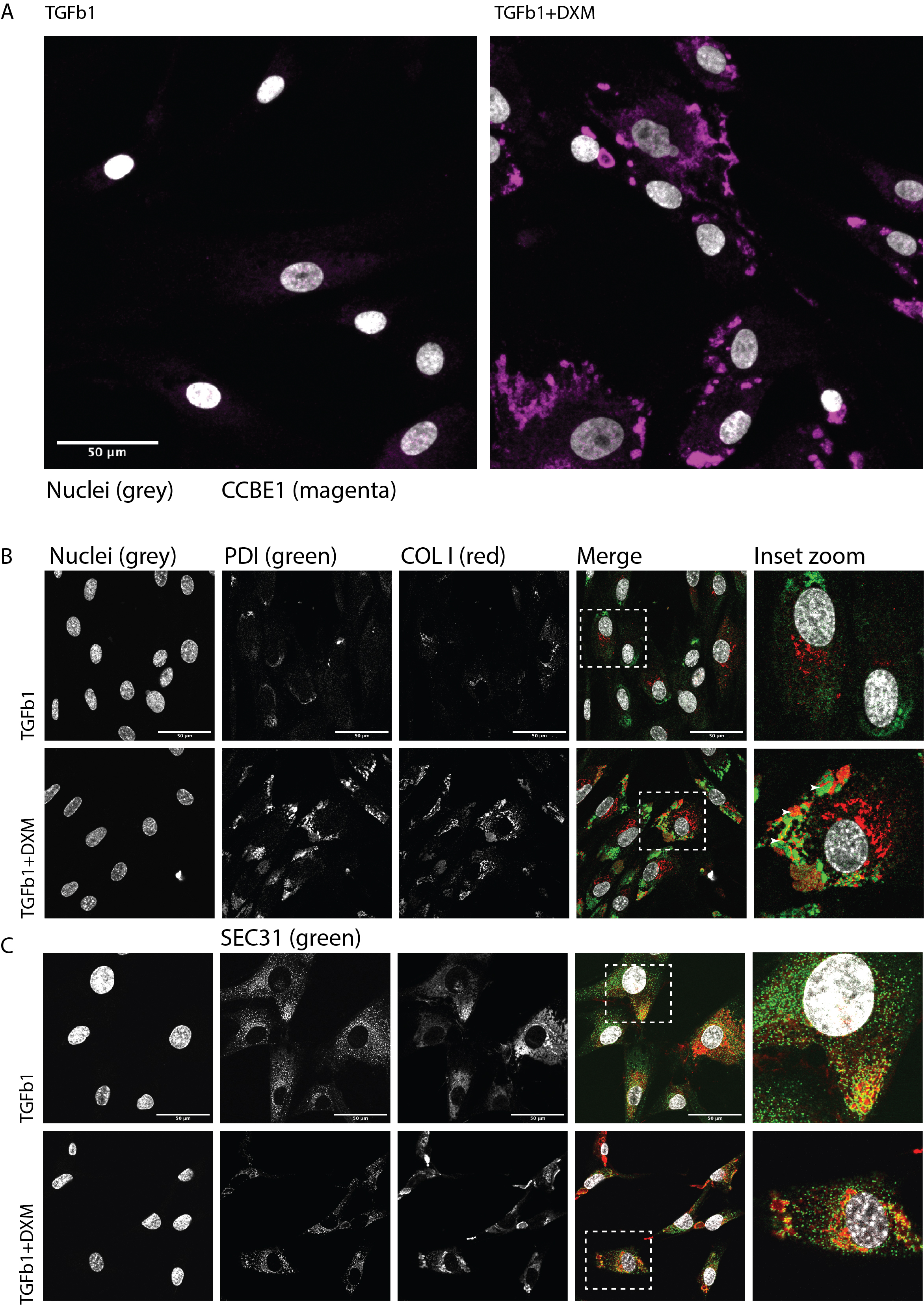

### Figure S3

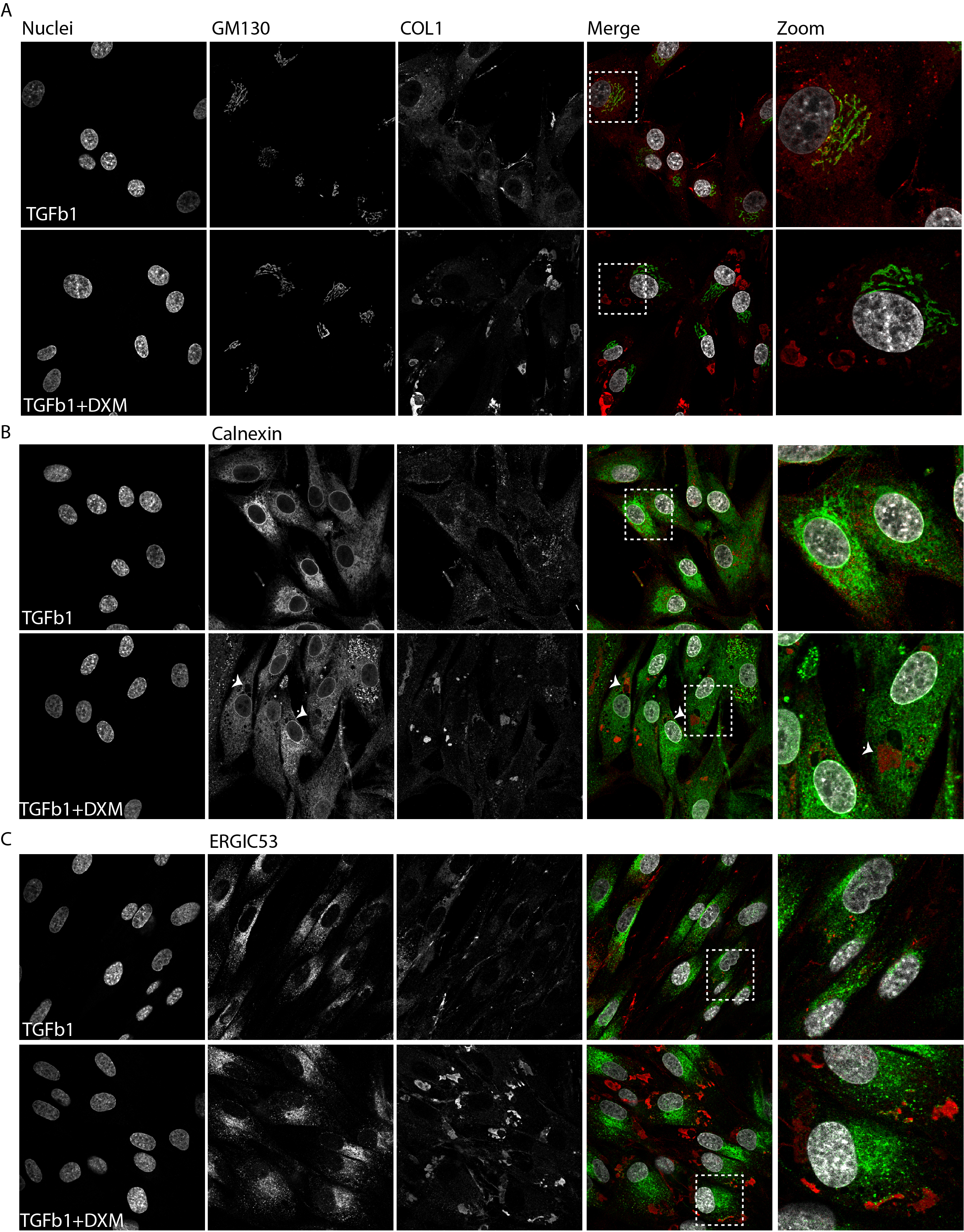

### Figure S4

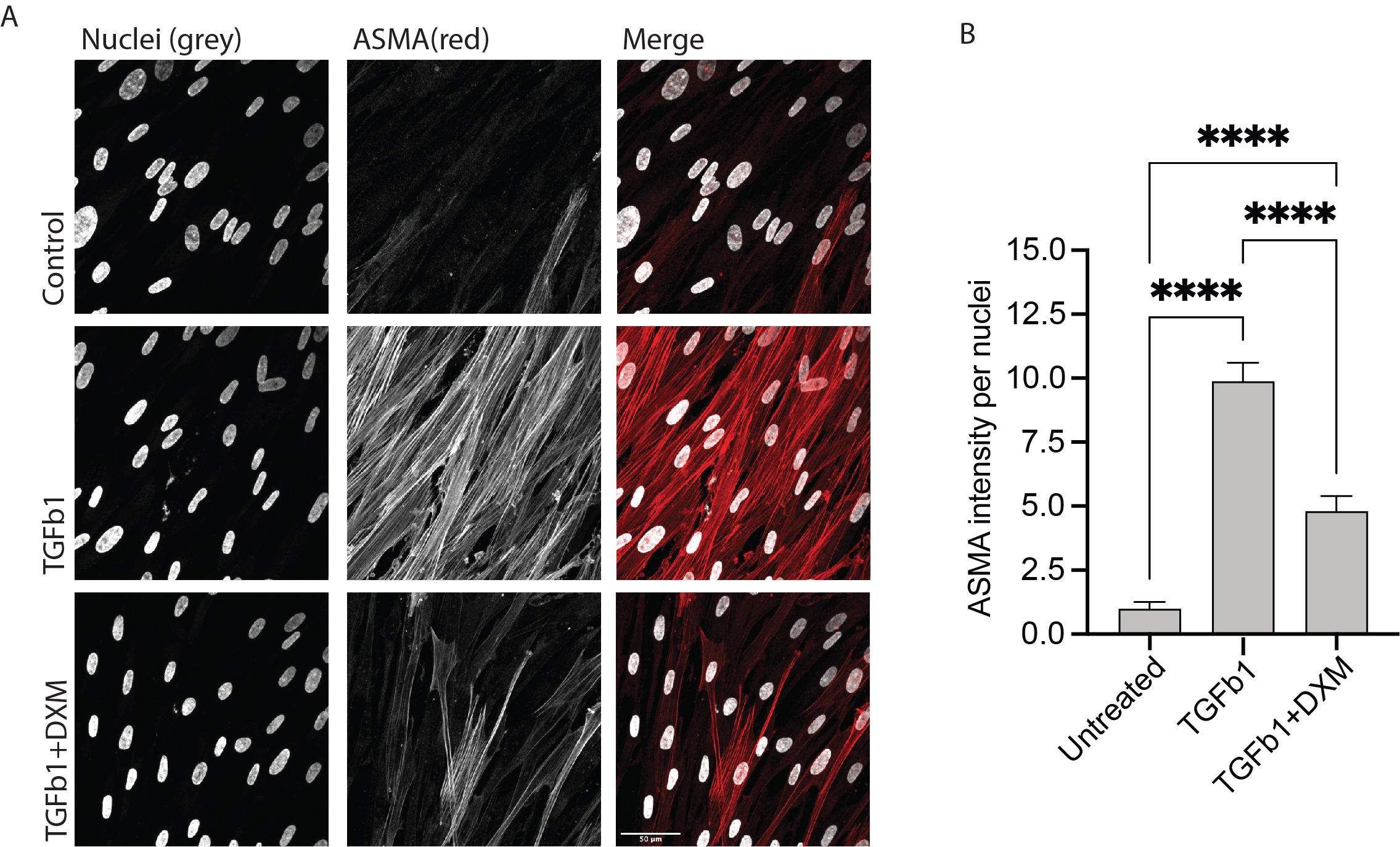
